# MUDflow: Combining Neural Networks, UMAP and DBM Clustering to Identify Cell Populations Accurately, Quickly and Easily in Mass and Fluorescence Cytometry

**DOI:** 10.1101/2025.06.19.660626

**Authors:** Connor Meehan, Stephen Meehan, Jonathan Ebrahimian, Wayne Moore, David Parks, Guenther Walther, Leonore A. Herzenberg

## Abstract

Identifying cell populations in flow cytometry data is mostly done via a “manual gating” method that often lacks verifiability and reproducibility, even in the hands of experienced investigators. Recently developed automatic gating methods have been shown to have good performance in cell population identification, but may require fine-tuned setup from experts or struggle to identify small populations. Here, we introduce an easily trainable multilayer perceptron neural network for automatic gating (MLPgater). Compared to the three popular automatic gating methods LDA, FlowSOM and PhenoGraph on three mass and six fluorescence cytometry datasets, MLPgater is most accurate by a substantial margin, tied with LDA as the fastest and uniquely able to replace manual gating except for training purposes. Furthermore, we show that combining MLPgater with UMAP’s guided dimensionality reduction feature and DBM’s clustering (MUDflow) effectively detects new populations that did not exist or were not identified in the training set.

## Introduction

Current flow cytometry instruments offer multi-laser measurements taken with detectors that capture entire emission spectra (“spectral” cytometers) or just a region around the peak emission of each dye (“conventional” cytometers). Mass cytometers use mass spectroscopy to measure isotope tracers as cell labels. Experiments may involve up to 50 dyes(1) or 60 mass isotopes(2), leading to formidable challenges in cell population identification and characterization, which is the foundation for much research discovery and many applications across numerous areas of medicine and biology.

Flow informatics aims to make data analysis reproducible, interpretable and investigator-independent. The field initially focussed on the collective visual presentation of cell measurements and introduced innovative algorithms such as probability contours, compensation and bi-exponential transformations. In all cases, the goal is/was to improve the investigator’s ability to see regions of interest within X/Y scatter plots and to draw boundaries, or “gates”, to delineate highly-populated regions (populations). Outlining such regions and subregions sequentially in this manner is known as “manual gating”.

Nevertheless, despite these innovations, manual gating remains imprecise, subjective and overly time-consuming(3,4). For example, experts frequently disagree on the placement of gates within a single bivariate plot, or on which sequence of data dimensions to use in composing a gating sequence. In 2006, bioinformaticians began to pursue fully automatic gating methods (AKA classifiers) for both research and clinical usages, particularly for high-dimensional data(5). As the number of measurements per cell has increased, so too has the need for at least one of these pursuits to succeed. One way to categorize these methods is as “supervised” or “unsupervised”. Supervised methods use a “training dataset” annotated with cell types in order to “learn” a classification. Unsupervised methods (often called “clustering” methods in flow cytometry) identify cell populations in cytometry data without any external information.

In 2015, Malek et al.(6) published such a method, named flowDensity, which outperformed state-of-the-art clustering algorithms by automatically replicating a predefined manual gating hierarchy based on assay-specific R code that processes density thresholds for every predefined gate. In 2018, Lux et al.(7) claimed that their newly-released method flowLearn “works the same way as flowDensity, but does not require a practitioner to manually tune hyper-parameters for an optimized outcome”. For each assay, however, flowLearn requires an expert in biology and density statistics to create gating hierarchies expressed as density thresholds in as many samples as flowLearn deems necessary. In 2019, Abdelaal et al.(8) published a supervised method using a linear discriminant analysis (LDA) classifier for mass cytometry data that achieves high F1-scores like flowLearn does but with much easier setup/training: for each assay, practitioners only input cell marker measurements and population labels from one or more representative samples. The population label can be from any prior gating that the practitioner regards as “ground truth”, manual or automatic.

LDA’s setup is similar to how one trains most neural networks for classification work. In 2007, Quinn et al.(9) first used neural networks with flow cytometry data for detecting cell apoptosis. In 2017, CellCNN(10) used them to predict outcomes with single-cell data. Then their effectiveness as generalized gating methods for mass cytometry data was established by DeepCyTOF(11) and DGCyTOF(12). In 2023, Suffian et al.(13) demonstrated accurate gating of conventional cytometry data with a 3-layer neural network.

Here we introduce a new automatic gating method (MLPgater) based on a neural network known as a “multilayer perceptron” (MLP). To maximize the usefulness of our automatic classifier, we strived to optimize its accuracy, speed, and ease of use in classifying data from conventional, mass and spectral cytometers. In our evaluations and method comparisons, we estimated the accuracy by F1-score or central similarity (see Methods) as compared to the reference gating. We measured speed by the total runtime of the classification method. We did not directly measure ease of use, but consider this goal achieved by MLPgater in that users only need to provide sample data and population labels to enable the analysis.

Manual gating is often used for new population discovery. By design, supervised methods only find known populations. On the other hand, unsupervised automatic methods are able to find both known and new populations, but infrequently use this distinction internally. For finding known populations, we can directly use the classification learned by MLPgater. Our solution for finding new populations (MUDflow) builds on the above classification as follows: the classification is used as “guidance” in a dimension reduction technique (UMAP(14)), after which we apply a density-based clustering method (DBM(15)), allowing us to identify additional populations among those previously classified as “background”. The combined result is a novel analysis pipeline that automatically delineates new populations.

MUDflow’s automation has the potential to replace manual gating for all scenarios except for the annotation of a training dataset. At CYTO 2023, flow informaticians estimated(16) that there were over 500 automatic methods, each of which is accurate in limited scenarios, thus echoing a conclusion of Lo et al. in 2008: “Like all clustering approaches, the methodology we have developed includes assumptions, which may limit the applicability of this approach, and it will not identify every cell population in every sample”(17). Our results show that MUDflow is highly accurate for many types of datasets and for very small populations, which provides a notable improvement in the applicability of automatic methods.

## Methods

In a flow cytometry assay, the researcher has a number *N* of defining cell markers (in this paper, *N* ranges from 8 to 43) that are used to make a measurement for each cell. Hence we can regard a flow cytometry cell measurement as a point *p* ϵ ℝ*^N^* The problem of cell population identification is to define a “classification”, which can be thought of as splitting ℝ*^N^* into *k* distinct subregions *S*_1_, …, *S_k_* This divides a flow cytometry dataset *B* into “cell populations”, each of which is a set of points lying within some subregion.

For a supervised method such as MLPgater, we must also have a training dataset *A* and a prior classification that gives a proper example of how to classify a sample. Typically, the prior classification is derived from manual gating (see Introduction). For the method to be successful, the test set *B* must be compatible with *A* in the sense that most or all of the cell phenotypes in the test set correspond to phenotypes in the training set, the *N* cell marker specificities and their dye or mass labels are identical and the measurements were taken under matched conditions (e.g., with the same settings on the same flow cytometer).

For situations like multi-site studies, we would hope that good standardization data from different sites and different instrument models would provide sufficiently “matched conditions”.

### Software and architecture choices

In deep learning, the ideal choice of architecture and hyperparameters for a neural network is highly dependent on the application and must usually be found experimentally by the practitioner(18). Through extensive testing, we settled on the following MLP architecture: a 5-layer fully-connected feedforward neural network with 3 hidden layers of 100, 50 and 25 neurons. Standard scaling is applied to all input data (we input *z* = (*x* − *u*)/*s*, where *x* is the observation and *u* and *s* are the mean and standard deviation, respectively, over the training set). To avoid overfitting, we use a dropout operation(19) in our 100-neuron layer that randomly does not apply 25% of the updates. For activations on the hidden layers, we use the rectified linear unit (ReLU)(20). For the output layer, we used softmax activation for classification(18). The optimization algorithm that we use for training is programming language-dependent; for MATLAB, we use a limited-memory BFGS algorithm(21); for Python, we use an Adam optimizer(22).

In our testing, we use both Python’s TensorFlow and MATLAB’s fitcnet. We settled on this architecture by comparing the results to those obtained using fitcnet’s optimization capability, which tests many combinations of hyperparameters and data partitions. We find that results from fitcnet, TensorFlow and this optimization of fitcnet are equivalent within the standard error of estimate.

To accelerate the computations, we rewrote UMAP(23) to be more suitable for tasks within our pipeline, such as guidance by MLPgater classifications. In addition, we identified default hyperparameter settings for UMAP and DBM(15) that are generally effective for flow cytometry data. For UMAP, we set the “reduced dimensionality” to 2, “nearest neighbors” to 15, the metric to “Euclidean”, the “minimum distance” to 0.3, and the “spread” to 1. We use “guided UMAP” (originally named “supervised UMAP”(24)), which reduces the weight of an edge in UMAP’s nearest-neighbors graph if it connects data points assigned to different populations. For DBM, we follow the guidelines in the supplementary materials of Meehan et al.(25) and set the “grid size” to 256, the “background scaling factor” to 4.3, and the “bandwidth” to 1.6.

Our software is designed to integrate directly with FlowJo 10.x. For flow analyses already stored in FlowJo, this enables the user to import features of the data setup (e.g., scaling, compensation, etc.), data measurements and labels of population definitions (both manual and automatic). This also enables convenient exporting of population definitions from MUDflow.

### MUDflow classification pipeline

#### Finding known populations

Classifying known populations with MLPgater can be thought of as a two-step procedure as explained below:

1. Using the training dataset *A* and pre-existing classification *C_A_*, train the neural network with architecture described in the previous section with the specified optimization algorithm. At completion, this yields a fixed network that assigns any point *p* ϵ ℝ*^n^* as belonging to a specific subregion, which mathematically can be expressed as a function *f*: ℝ*^N^* → {0, 1, 2, …, *k*}. Here, *f*(*p*) = 0 if *p* is considered “background” (e.g., a doublet cell, debris, or otherwise not belonging to the defined populations).
2. Use the network to create a classification *C_B_* for the test dataset *B* by partitioning ℝ*^N^* so that every cell *p* in *B* lies in the subregion associated with *f*(*p*).

The details of the neural network training in step 1 are rather complex, but the training follows a process that is fairly standard in the domain of machine learning. Each neuron in the network accepts a vector-valued input x and outputs the ReLU function *max*{0,**w** · **x** + *c*}, where the vector w and the scalar *c* are parameters that can be adjusted during training. The final vector output z is transformed via the softmax function (softmax(**z**)*_i_* =exp(*z_i_*)/∑*_j_* exp(*z_j_*)) and compared to the original classification *C_A_* via cross-entropy. More details can be found in most standard references on neural networks (e.g., Goodfellow et al.(18)).

#### Finding new populations

When using MUDflow to find new populations, we perform the following three steps in addition to the two from the previous section:

3. Perform dimensionality reduction on the test dataset *B* with uniform manifold approximation and projection (UMAP)(14) to create a two-dimensional “embedding” *E : B* → ℝ^2^. Specifically, we perform “guided UMAP” using the classification *C_B_* from step 2 as the guide.
4. Use density-based merging (DBM) clustering(15) directly on the dimension-reduced dataset *E*[*B*], which induces another classification *C_MUD_* that divides *B* into *m* clusters. Typically, *m*>*k*.
5. Use QFMatch(26) to create a matching *Q* of populations of *B* defined by the classifications *C_B_* (MLPgater populations) and *C_MUD_* (clusters defined by DBM). If a cluster is not matched to any MLPgater population by *Q*, we denote it as a “new population”. If *Q* matches multiple clusters to a single MLPgater population, we denote the clusters as “new subpopulations”.

Hence MUDflow, as a method, can be considered a proper extension of MLPgater. Henceforth, we will call our method “MLPgater” when we are only considering the first two steps and “MUDflow” otherwise. See Figure 1 for a schematic diagram of the entire pipeline.

**Figure 1.**
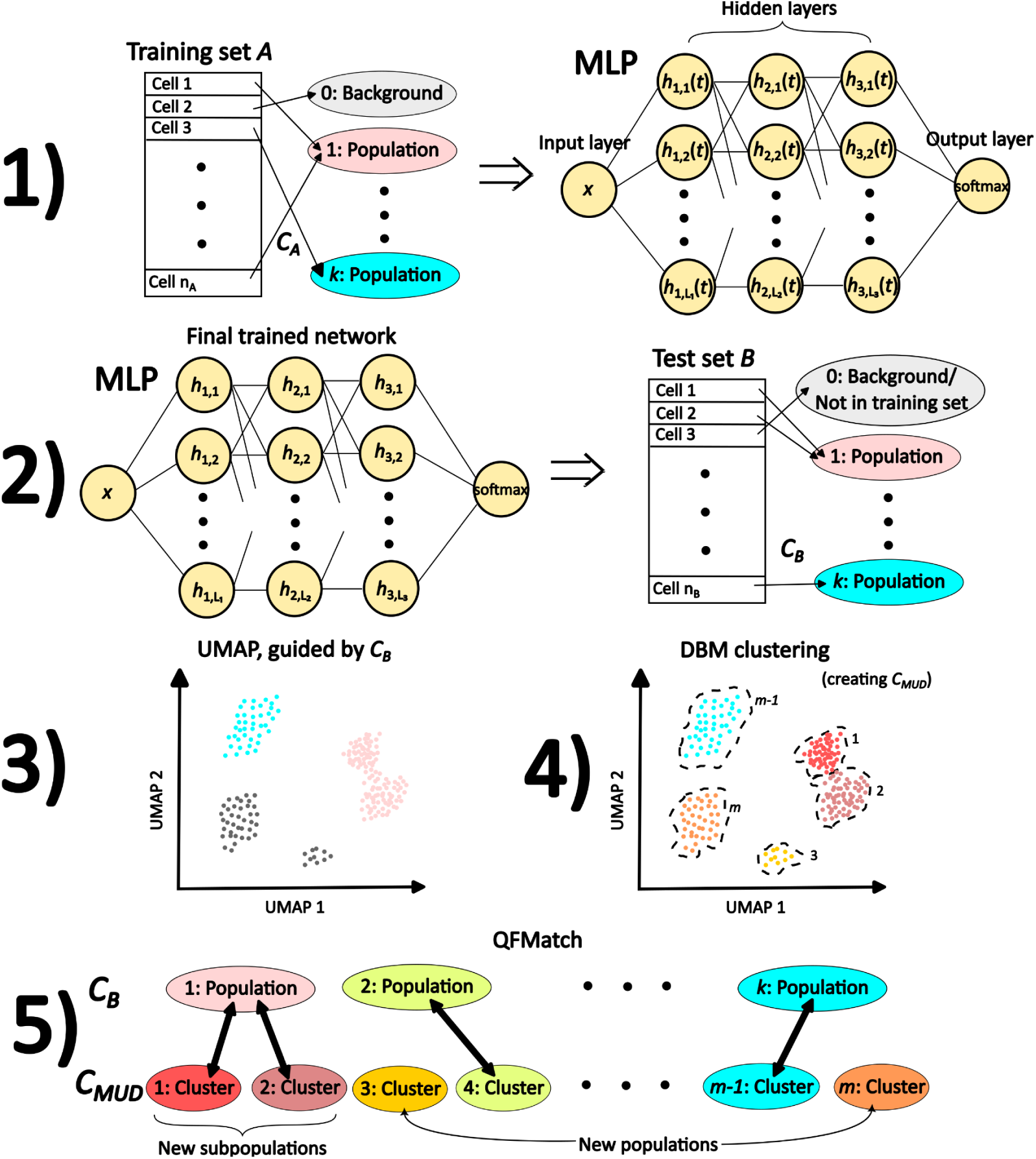
Overview of the MUDflow classification pipeline. 1) A fully-connected feedforward neural network (MLP) is trained with a training dataset *A* and a pre-existing classification *C_A_* (typically created through manual gating). 2) The network is used to create a classification *C_B_* for a new, unannotated test dataset *B*. 3) For new population discovery, the pipeline is continued by creating a two-dimensional representation of *B* using guided UMAP with *C_B_* used as the guide. 4) DBM is used to define *C_MUD_*, which labels clusters found in the UMAP plot. 5) QFMatch is used to create a matching between populations found by the neural network and clusters found by DBM in the UMAP plot. Some of the DBM clusters can then be called “new populations” and “new subpopulations”.

### Classifier evaluations and comparisons

The inventors of both flowDensity(6) and flowLearn(7) demonstrated the accuracy of their methods by comparing their results to other established supervised (DeepCyTOF(11)) and unsupervised methods (FlowSOM(27), X-Cyt(28), SamSpectral(29) and flowMeans(30)). In a similar approach, here we compare MLPgater to LDA(8) (a supervised method), FlowSOM and PhenoGraph(31) (unsupervised methods), which Liu et al. ranked as the three best methods overall in 2019(32).

### Datasets description

We tested our pipeline on nine published datasets, treating the reported peer-reviewed manual gating analyses as ground truth. For more details, see Table 1 and Supplementary Section S1.

**Table 1.** The names of nine published datasets used for testing in this paper, as well as the number of populations and defining markers (parameters) associated with each. We omitted 3 populations from the OMIP-069 study (noted with an asterisk) because they had substantial overlap with the other populations.

| Publication | Cytometer type | Number of samples used | Number of folds for cross-validation | Number of defining markers | Number of populations reported |
| --- | --- | --- | --- | --- | --- |
| OMIP-044(45) | Conventional | 3 | 3 | 14 | 13 |
| OMIP-047(46) | Conventional | 6 | 4 | 9 | 8 |
| OMIP-058(34) | Conventional | 2 | 2 | 17 | 35 |
| OMIP-069(39) | Spectral | 4 | 3 | 26 | 43* |
| OMIP-077(38) | Conventional | 4 | 4 | 14 | 14 |
| ESHGHI(37) | Mass | 10 | 5 | 27 | 28 |
| GHOSN(36) | Conventional | 3 | 2 | 9 | 13 |
| LEIPOLD(47) | Mass | 6 | 4 | 15 | 18 |
| PANORAMA(41) | Mass | 10 | 5 | 25 | 24 |

### Performance metrics

Here, we use two performance metrics to gauge the success of the automatic classifiers tested. The first is F1-score, which measures population overlap via the harmonic mean of precision and recall. Since FlowCAP(33), most flow informatics publications assess gating accuracy using the F1-score.

Secondly, we use Central similarity (CS), which measures overlap in the central dense region of combined populations with the Jaccard index. We remove the 20% of the cells in the union of the populations that are farthest from the union’s mean using Mahalanobis distance. This definition is based on a key characteristic of phenotypes: their event measurements show a marked central tendency in all dimensions, indicating that they are “atomic”, i.e., not splittable in all tested dimensions.

### Testing procedures

#### Finding known populations

To evaluate the performance of the four classifiers, we run the following four-step procedure for each of the nine datasets.

1. Prepare the data using FlowJo 10.10. We use the *logicle* transform to reproduce the published data scales (see Table 1) illustrated in the manual gating in the corresponding original paper. We use the embedded compensation matrix for conventional cytometry data (except with OMIP-058(34), for which we computed it using FlowJo’s compensation wizard).
2. Manually reproduce the published manual gating hierarchy for non-overlapping “leaf gates” (gates that have no subgates as children). Seven of 204 published leaf gates are excluded because they have many cells in common with other leaf gates.
3. Perform automatic gating on the smallest gate containing all leaf gates, using all associated data for those parameters which define the branch’s subgates (subgate-defining parameters). Crucially, this is distinct from the procedure of Abdelaal et al. for testing LDA(8), in which they discard cells that are not in leaf gates (see Supplementary Section S3). Since the membership of cells in leaf gates is not typically known *a priori*, we feel it is more realistic to include all cells.

a. Run MLPgater and LDA (with their MATLAB implementations) using *M*-fold cross-validation, a common technique for evaluating the prediction error of a machine learning model(35). The full dataset is partitioned into *M* similarly-sized groups of samples while leaving each biological sample fully intact within exactly one of the *M* groups. For each group in turn, it is used as the test set for the performance metrics, while the remaining groups act as the training set. The value of *M* varies by dataset (see Table 1).
b. Run FlowSOM and PhenoGraph using the plugins from FlowJo 10.10. For FlowSOM, we set hyperparameters as described in Supplementary Section S2. For PhenoGraph, we use 30 for its only hyperparameter “K” (nearest neighbors).
c. Match the unlabelled FlowSOM and PhenoGraph populations one-to-many with manually gated populations using QFMatch, which merges subsets to find the best F1-score (see Orlova et al.(26), step 4 on page 5 and Figure 3 on page 6).
4. Store automatic gates in a workspace file readable by FlowJo or other software (e.g., flowWorkspace from Bioconductor).

**Figure 3.**
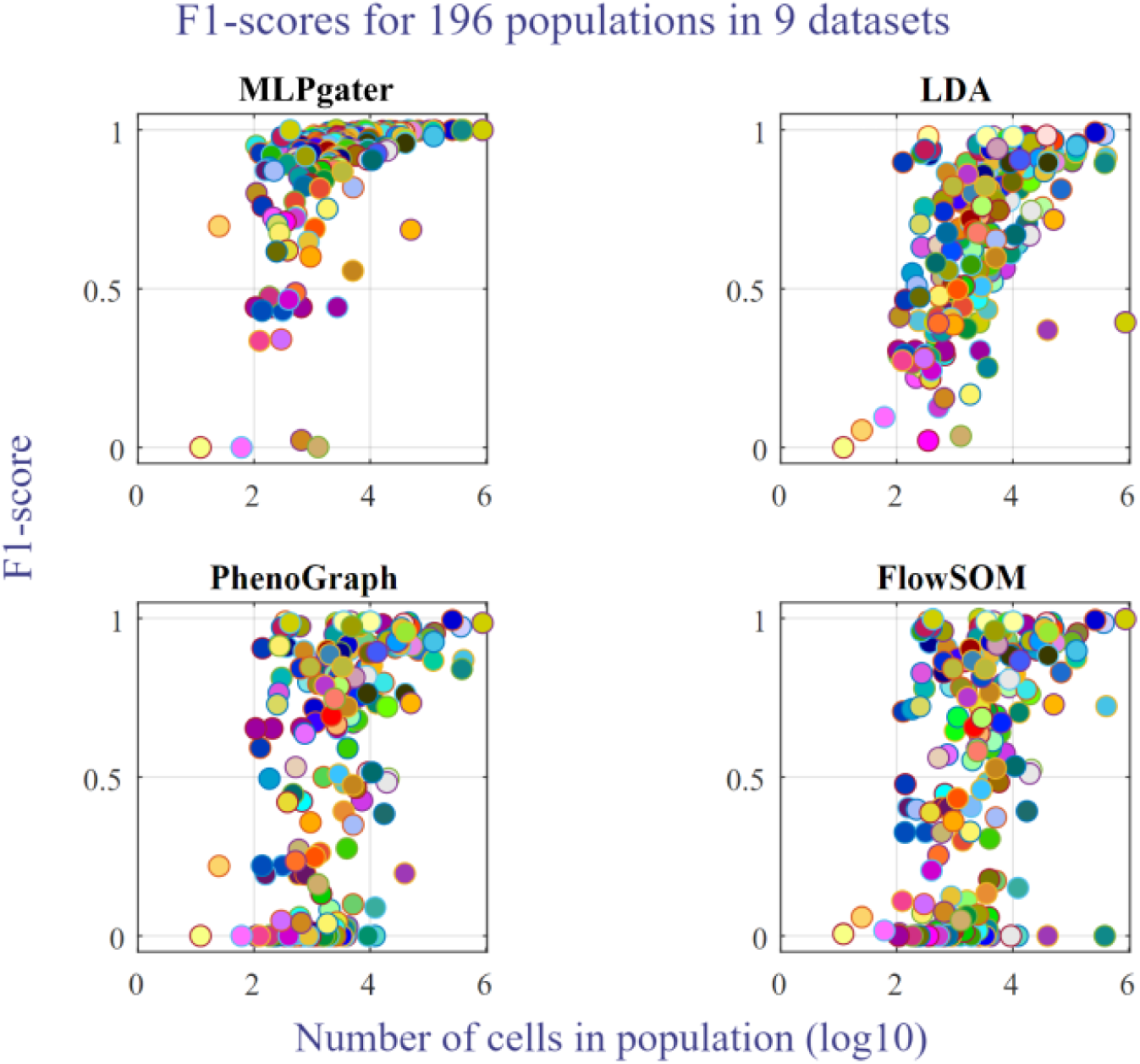
Classifier performance relative to population size. For each of the four classifiers, a scatter plot shows the F1-score achieved by the classifier on a population vs. the population’s size on a logarithmic scale. Smaller populations tend to be harder for automatic classifiers to locate. See Supplementary Table S18 for a legend of population colors.

#### Finding new populations

We investigate the identification of new populations only for the GHOSN(36) and ESHGHI(37) datasets. We first prepare a special training dataset following steps 1 and 2 above. We then follow the five steps presented in the section “MUDflow classification pipeline”, using the performance metrics to evaluate the resulting classification.

With the GHOSN dataset, we train MLPgater on a sample from a “RAG knockout” mouse strain that is genetically modified to not produce lymphocytes. The resulting network is used to identify populations in samples from a BALB/c mouse that has lymphocytes. With the ESHGHI dataset, we train MLPgater on a sample from which we have removed the published manual gates for eosinophils, neutrophils, basophils, plasmablasts, mDCs and pDCs.

The new population data are present in the ESHGHI training as background. Thus, the GHOSN training is more representative of a real-world use case, since the new population data are genetically knocked out and thus cannot bias the training.

## Results

### Finding known populations

For each of the 9 tested datasets, Figure 2 shows the F1-score distribution across populations and Table 2 shows the median and mean for the F1-score and CS.

**Figure 2.**
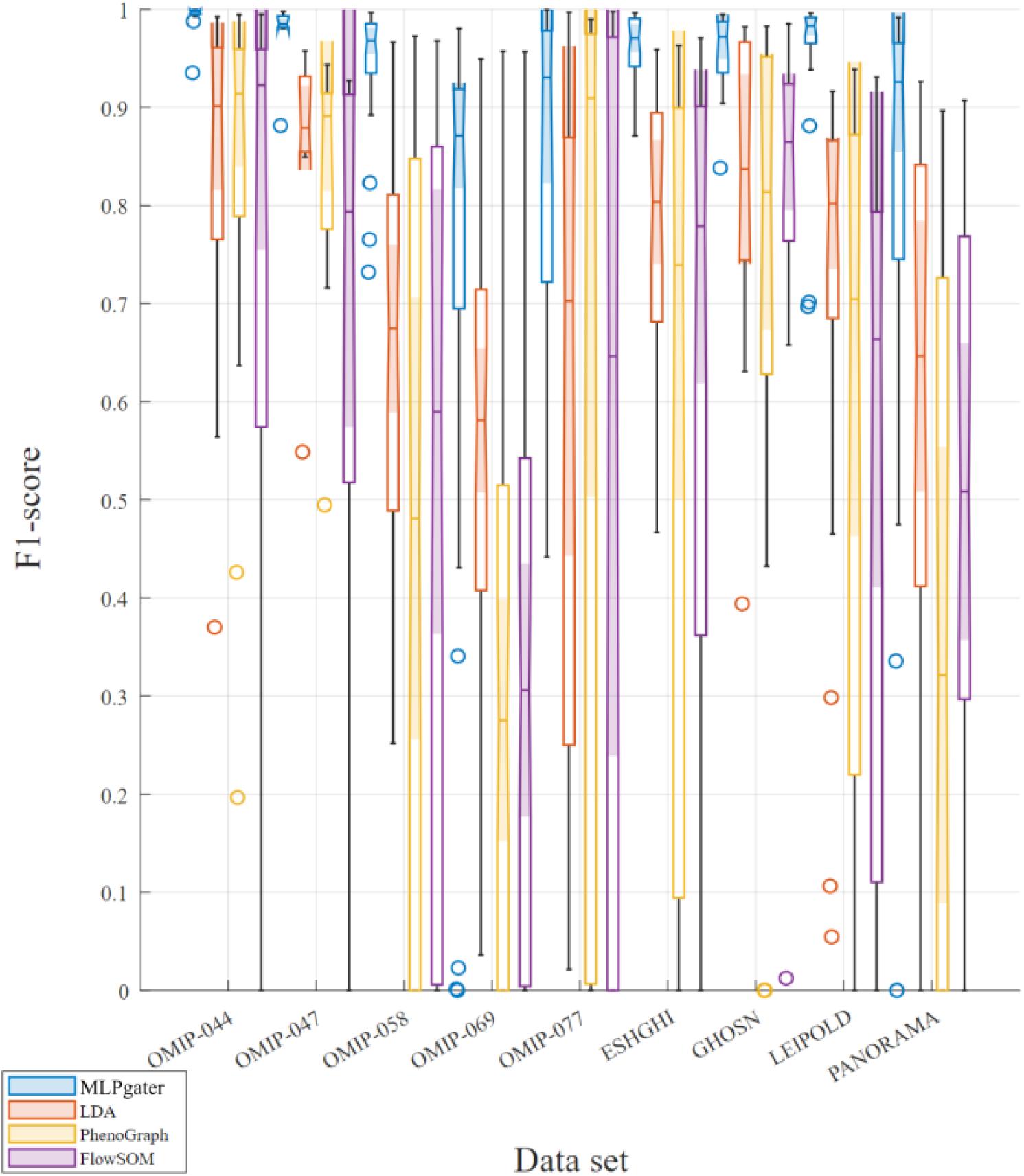
Classifier performance comparison. For each of the nine datasets and four classifiers discussed in Methods, a box-and-whisker plot shows the median and quartiles of the F1-scores for the dataset’s populations, as determined by the original publication’s manual gating. Outliers are computed with interquartile range and denoted by circles. MLPgater has the highest median score for each dataset.

**Table 2.**
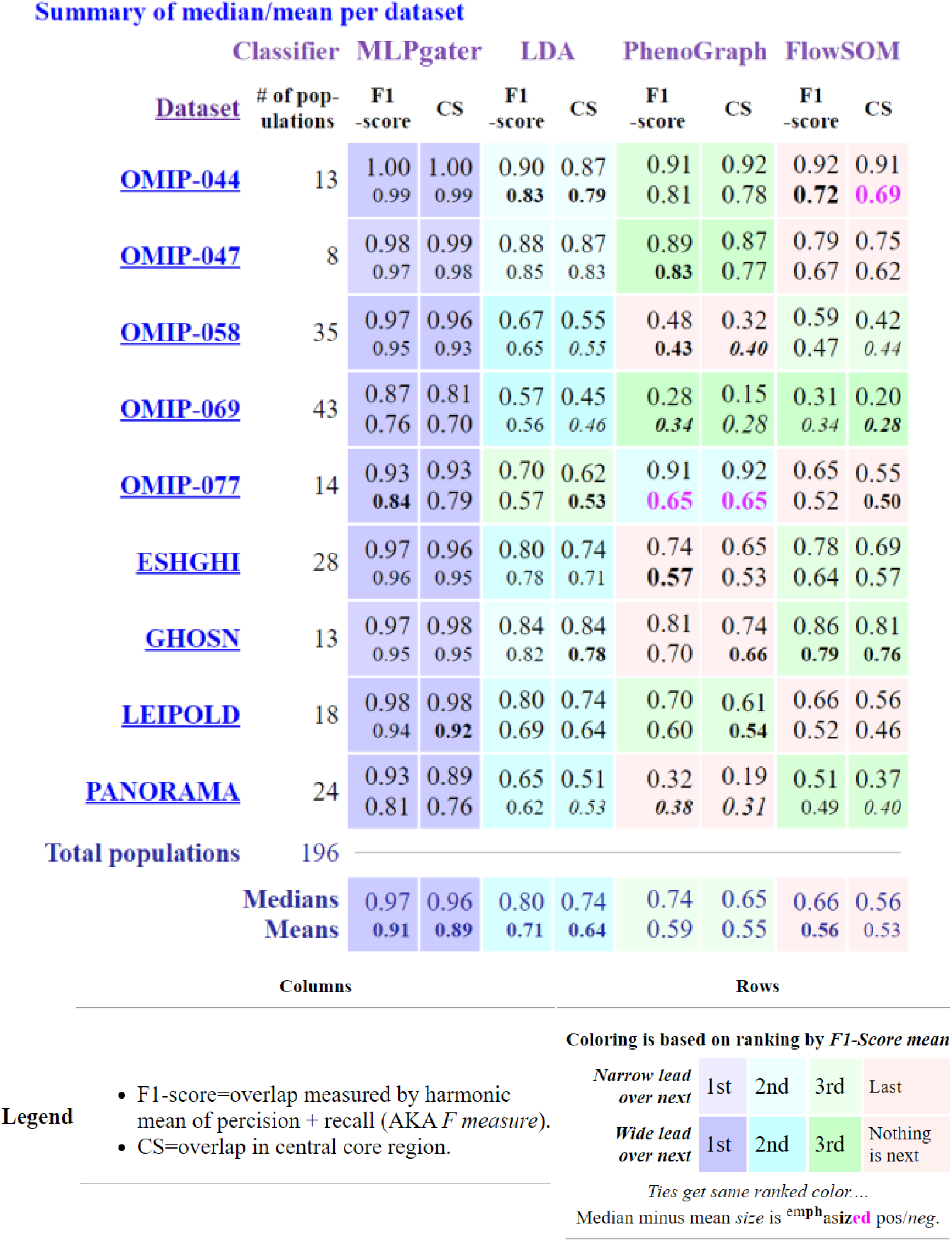
Summary values of F1-score and central similarity (CS) for each classifier and dataset. In each cell, both the median and mean (over all of the populations in each dataset) value of F1-score and CS are given. Cells are ranked by median F1-score and colored accordingly. The overall medians rank MLPgater as the best classifier, followed by LDA, PhenoGraph, and FlowSOM, respectively.

MLPgater has the highest scores for F1-score and CS. Furthermore, it has the least variation between the F1-score and CS and between the medians and means, and the least variation among the datasets for the same performance metric. The CS difference between MLPgater and the other classifiers is greater than the F1-score difference in all cases except with OMIP-077(38). This indicates that MLPgater is particularly sensitive and suited to the needs of phenotype classification.

The accuracy of LDA, PhenoGraph and FlowSOM varies substantially between datasets even when the matched manual gates are equally plausible-looking. In contrast, MLPgater’s accuracy is more consistent and robust despite dataset differences.

The F1-score and CS per population can be seen in Supplementary Section S9. Supplementary Section S6 discusses how OMIP-069’s(39) lower scoring is partly due to manual gating issues specific to this publication.

#### Population size issues

The F1-scores versus population size for the four methods tested are shown in Figure 3.

Automatic methods often miss smaller populations(40). MLPgater yields F1-scores of at least 0.70 in all 115 populations of six datasets, as illustrated in Figure 4.

**Figure 4.**
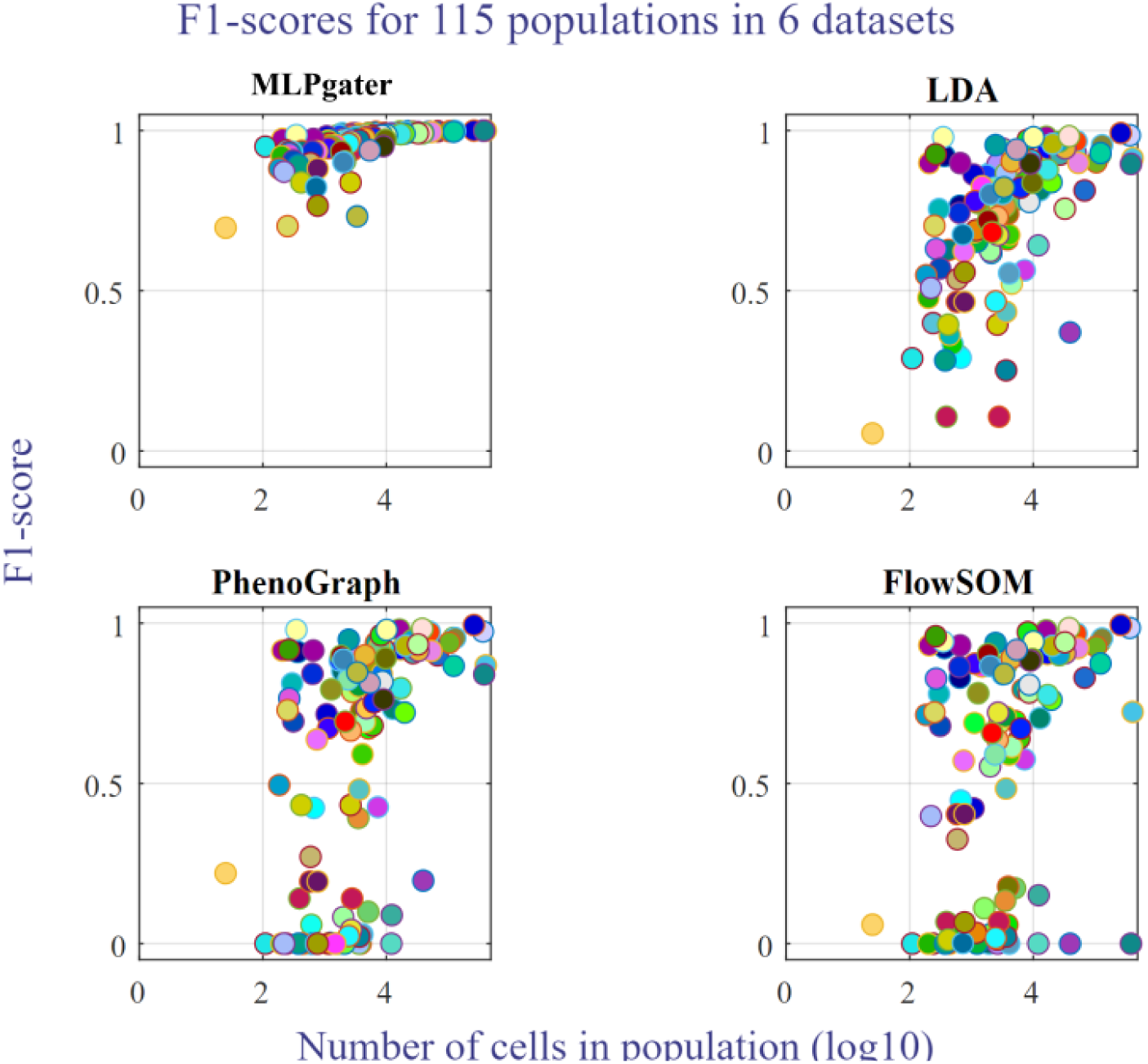
The same scatter plots as in Figure 3, restricted to the six of nine datasets on which MLPgater achieves no F1-scores below 0.50. See Supplementary Table S18 for a legend of population colors.

In the three other datasets, 11 of 81 populations have MLPgater F1-scores less than 0.50: one in OMIP-077 (0.44), four in PANORAMA(41) and six in OMIP-069. Table 3 explores the performance of classifiers on populations with the lowest F1-scores. For more details, see Supplementary Section S5.

**Table 3.** Difficulty with small populations. For each of the nine datasets and four classifiers discussed in Methods, the table shows the smallest F1-score achieved, the number of populations for which an F1-score of less than 0.50 was achieved, and the median frequency of these populations.

|  | Minimum F1-score |  |  |  | Number of populations less than 0.5 F1-score |  |  |  | Median frequency of populations less than 0.5 F1-score (%) |  |  |  |
| --- | --- | --- | --- | --- | --- | --- | --- | --- | --- | --- | --- | --- |
| Dataset | MLP-gater | LDA | Pheno-Graph | FlowSOM | MLP-gater | LDA | Pheno-Graph | FlowSOM | MLP-gater | LDA | Pheno-Graph | FlowSOM |
| OMIP-044(45) | 0.94 | 0.37 | 0.20 | 0 | 0/13 | 1/13 | 2/13 | 2/13 |  | 2.0 | 1.2 | 11 |
| OMIP-047(46) | 0.88 | 0.55 | 0.49 | 0 | 0/8 | 0/8 | 1/8 | 2/8 |  |  | 0.38 | 3.9 |
| OMIP-058(34) | 0.88 | 0.25 | 0 | 0 | 0/35 | 9/35 | 6/35 | 11/35 |  | 0.12 | 0.77 | 0.20 |
| OMIP-069(39) | 0 | 0.04 | 0.04 | <0.01 | 6/43 | 16/43 | 16/43 | 18/43 | 0.08 | 0.13 | 0.33 | 0.26 |
| OMIP-077(38) | 0.44 | 0.02 | 0 | 0 | 1/14 | 6/14 | 2/14 | 4/14 | 0.01 | 0.03 | 0.04 | 0.02 |
| ESHGHI(37) | 0.87 | 0 | 0 | 0.82 | 0/28 | 8/28 | 8/28 | 0/28 |  | 1.1 | 1.1 |  |
| GHOSN(36) | 0.77 | 0.39 | 0.43 | 0.01 | 0/13 | 1/13 | 1/13 | 1/13 |  | 0.36 | 0.36 | 0.36 |
| LEIPOLD(47) | 0.65 | 0.05 | 0 | 0 | 0/18 | 4/18 | 6/18 | 7/18 |  | 0.89 | 1.2 | 1.4 |
| PANORAMA(41) | 0 | 0 | 0.01 | 0 | 4/24 | 8/24 | 7/24 | 12/24 | 0.30 | 0.68 | 1.8 | 1.1 |
| **Totals** |  |  |  |  | 11/196 | 47/196 | 49/196 | 65/196 |  |  |  |  |

As with most machine learning methods, using a larger training dataset tends to result in better performance with MLPgater. The smallest training set we used was about 125,000 cells (GHOSN dataset) on which MLPgater performed quite well (minimum F1-score of 0.77, median score of 0.95). However, in general, we recommend that the training set have size at least two times the sample size that an assay designer deems as the minimum feasible for a particular assay; larger if some populations have less than 1% frequency.

#### Improving accuracy

As discussed in step 3a of “Testing procedures”, we train with reference cell populations and analyze new cell samples by including all of the cells associated with a branch gate in the computation. In contrast, Abdelaal et al.(8) train and identify using only the cells that occur in the leaves of the branch gate. Our broader testing shows that such filtering actually improves LDA’s scores more than it improves MLPgater’s (see Supplementary Tables S1 and S2). In any event, since the goal is to replace manual gating, it is unreasonable to base classifications on pregated data.

FlowSOM(27) and PhenoGraph(31) accuracy improves when testing with many-to-many matching (instead of the one-to-many matching in step 3c of “Testing procedures”), which reveals good matches between one or more automatic populations and one or more manually gated populations (see Supplementary Tables S3 and S4). Nevertheless, because we treat scientist-reviewed manually gated populations as ground truth, we do not allow manual gates to be merged in the matching process in our normal testing procedure. The accuracy also improves on smaller subregions of a publication’s manual gating hierarchy with fewer subgate-defining parameters. For example, PhenoGraph and FlowSOM find the same populations more accurately when starting with 9 parameters at the parent gate of T cells and NKT-like cells than when starting with 26 parameters at the gate common to all OMIP-069 populations; see Supplementary Tables S7 and S8. In contrast, MLPgater’s scores are far less affected by altered starting points even when starting at the sample level which includes doublets, debris, dead cells and off-scale measurements; see Supplementary Section S7.

Score improvements also occur when, for a particular classifier and dataset pairing, we add specific R/MATLAB/Python programming and statistical tuning like flowDensity requires, albeit for each and every gate. While our implementation of MLPgater allows for this, such customization conflicts with one of our three primary automation goals: ease of use.

#### Classification speed

We measured the runtime of the four classifiers on the GHOSN dataset using a computer with 6 cores and 32 GB RAM. For training the model of supervised methods, we used a training set with 126,539 cells, each with 9 parameters, from the peritoneal cavity of a BALB/c mouse (with file name “all_3-3.fcs”). Training a classifier model from this dataset requires 67 seconds for MLPgater and 0.48 seconds for LDA. Training a prior map for FlowSOM (not used in this paper) requires 17 seconds.

To measure the runtime of the classification, we used a test set with 133,569 cells and 9 parameters from the peritoneal cavity of a C57BL/6 mouse (with file name “all_3-1.fcs”). The runtime was 0.11 seconds for MLPgater, 0.13 seconds for LDA, 298 seconds for PhenoGraph, and 5 seconds for FlowSOM.

Training for MLPgater and LDA occurs when initially setting up an assay. Thereafter, that assay rarely requires retraining (provided the original training is truly representative), even when adding new markers for further discovery. Importantly, the computation time of LDA and MLPgater increases linearly as sample size increases.

### Finding new populations

When run on the GHOSN dataset with mouse peritoneal cavity cells, step 3 of our method presented in “MUDflow classification pipeline” produces the embedding in two-dimensional space shown in Figure 5. The purple curves represent the boundaries of clusters found by step 4.

**Figure 5.**
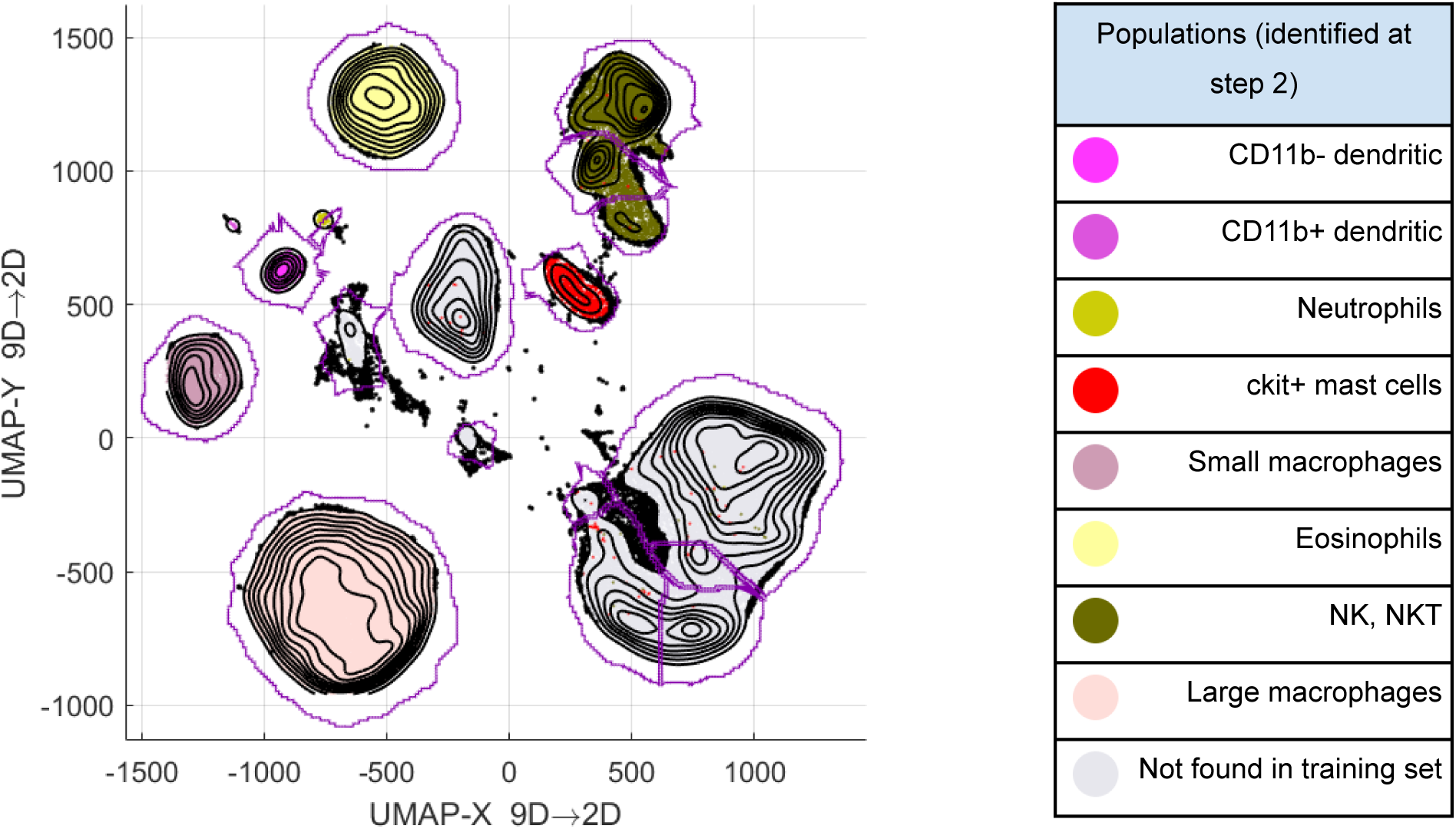
Guided UMAP plot of flow cytometry data from a BALB/c mouse peritoneal cavity (PerC). The classification used for guidance was generated by MLPgater trained on RAG mouse data. The BALB/c mouse sample contains lymphocytes not seen in the RAG mouse sample, leading to the large uncolored groups of data labeled “Not in training set”.

The colored clusters in Figure 5 contain populations that MLPgater identified in a normal sample (BALB/c strain) from training with a sample that lacks lymphocytes (RAG strain). Using our software to highlight populations from the sample’s manual gating shows that the lymphocytes occur mostly in the uncolored clusters. The smallest population, “Developing B cells”, is 1,000 cells or 1.3% of the total cells in the new populations found. Figure 6 and Table 4 show how closely the new populations found by our method align with the manually gated populations.

**Figure 6.**
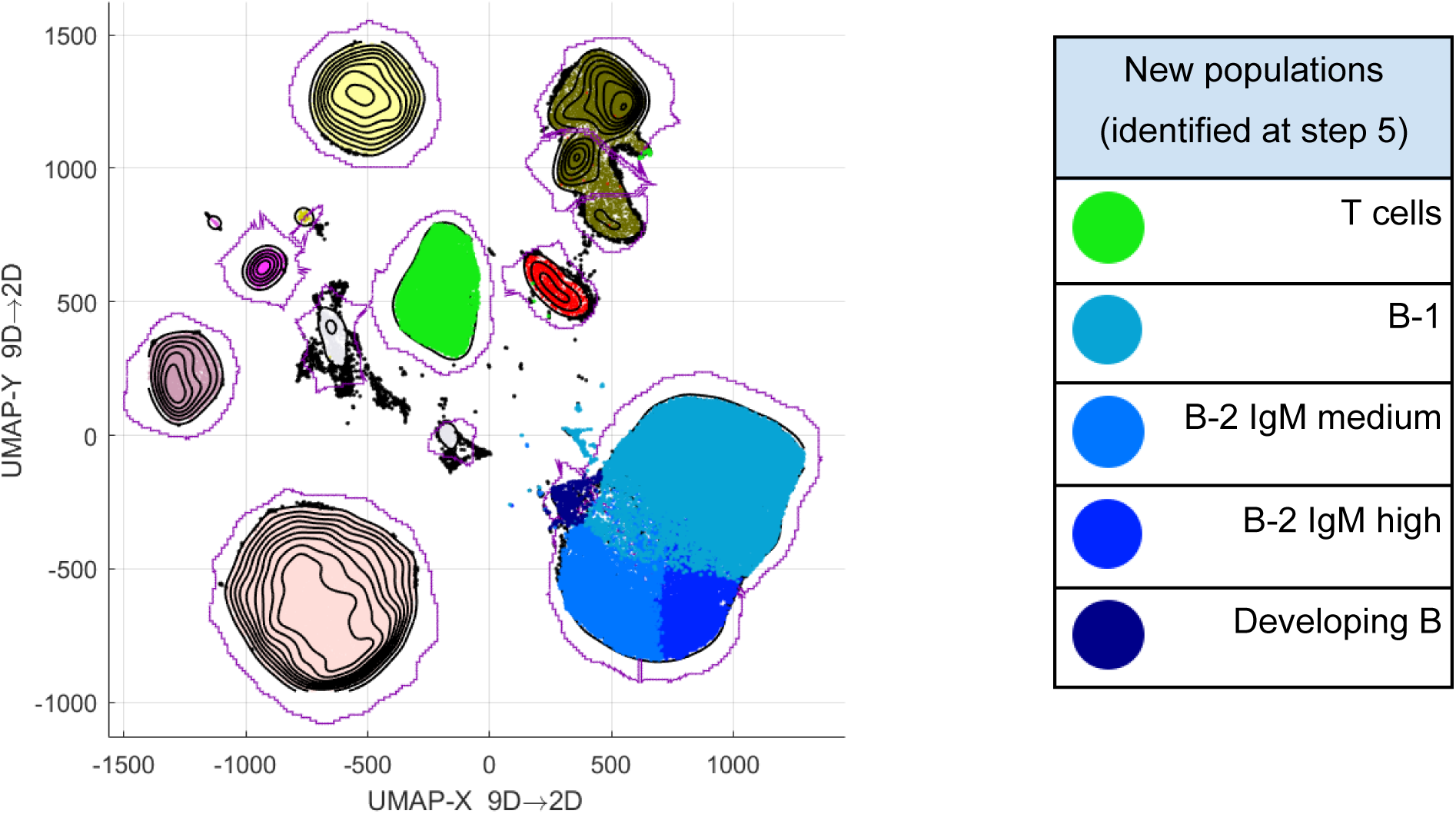
Guided UMAP plot from Figure 5, after (1) running DBM clustering and (2) using QFMatch to match DBM clusters one-to-many with manually gated populations from the GHOSN publication.

**Table 4.** Comparison of dimension reduction techniques for finding new populations, when used in step 3 of MUDflow, in the GHOSN and ESHGHI datasets. F1-scores and CS values are reported for new populations in the datasets using our method with the specified dimension reduction technique. New populations are biologically novel in the GHOSN dataset, whereas they have been manually removed from the ESHGHI dataset. An empty cell indicates that the technique did not find the corresponding population. Missed populations are assigned a value of 0 for the purpose of median and mean calculations.

| GHOSN dataset |  |  |  |  |  |  |  |
| --- | --- | --- | --- | --- | --- | --- | --- |
|  |  | Guided UMAP |  | Basic UMAP |  | PHATE |  |
| Population | Frequency (%) | F1-score | CS | F1-score | CS | F1-score | CS |
| T cells | 16 | 0.98 | 0.99 | 0.97 | 0.98 | 0.96 | 0.95 |
| B-1 | 53 | 0.92 | 0.92 | 0.92 | 0.92 | 0.91 | 0.91 |
| B-2 IgM medium | 17 | 0.84 | 0.81 | 0.84 | 0.81 | 0.72 | 0.63 |
| B-2 IgM high | 12 | 0.72 | 0.64 | 0.72 | 0.63 |  |  |
| Developing B | 1.3 | 0.85 | 0.77 |  |  |  |  |
| Median |  | 0.85 | 0.81 | 0.84 | 0.81 | 0.72 | 0.63 |
| Mean |  | 0.86 | 0.83 | 0.69 | 0.67 | 0.52 | 0.50 |
| ESHGHI dataset |  |  |  |  |  |  |  |
|  |  | Guided UMAP |  | Basic UMAP |  | PHATE |  |
| Population | Frequency (%) | F1-score | CS | F1-score | CS | F1-score | CS |
| Neutrophils | 18 | 0.96 | 0.96 | 0.96 | 0.96 | 0.94 | 0.92 |
| mDCs | 5.2 | 0.56 | 0.43 | 0.56 | 0.43 | 0.56 | 0.43 |
| Eosinophils | 15 | 0.90 | 0.84 | 0.90 | 0.84 |  |  |
| pDCs | 8 | 0.88 | 0.83 |  |  |  |  |
| Basophils | 52 | 0.84 | 0.79 |  |  |  |  |
| Plasmablasts | 2.4 | 0.76 | 0.72 |  |  |  |  |
| Median |  | 0.86 | 0.81 | 0.28 | 0.21 | 0 | 0 |
| Mean |  | 0.82 | 0.76 | 0.4 | 0.37 | 0.25 | 0.23 |

Figure 6 also shows the discovery of new subpopulations of the NK, NKT population identified by MUDflow.

Following the same procedure with the ESHGHI dataset, we get the F1-scores shown in Figure 7 and Table 4. The smallest population (plasmablasts) is 400 cells.

**Figure 7.**
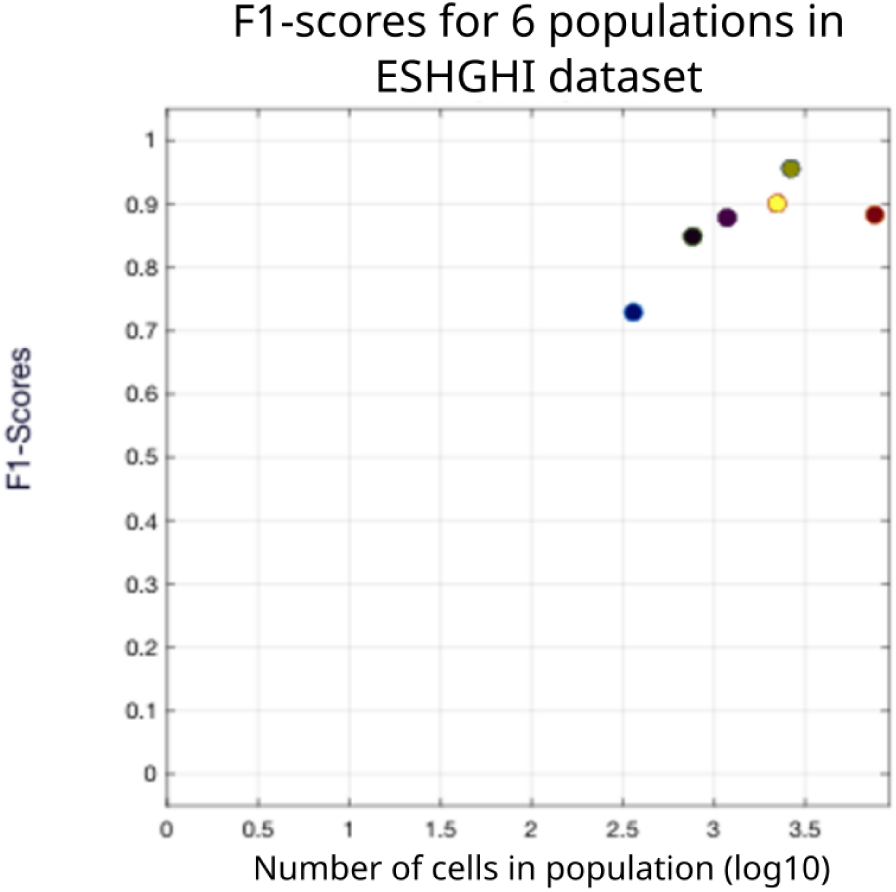
The F1-score of each new population from the ESHGHI dataset vs. the population’s size on a logarithmic scale. All 6 populations were not present in the training set for MUDflow, but found later using UMAP dimensionality reduction and DBM clustering.

From a theoretical standpoint, it seems equally valid to replace guided UMAP in step 3 of our new population discovery method with another dimension reduction technique. To justify our choice, we compare guided UMAP with two other forms of dimension reduction: basic UMAP(14) and PHATE(42). Basic UMAP is identical to guided UMAP, except that it does not receive classification of data points and hence skips the weight-adjusting step mentioned in the section “Software and architecture choices”. Note that directly comparing basic UMAP plots of different datasets comes with difficulty; see Supplementary Section S8. PHATE is a non-linear dimension reduction technique introduced by Moon et al. in 2019 that was found to represent global and local structure well for mass cytometry data sets(42).

Figure 8 and Table 4 below summarize the results of our testing. New population discovery is not as good when dimension reduction is not guided by MLPgater’s known population definitions. Future research will explore this with more datasets, other dimension reduction algorithms and experimentation of the algorithms’ hyperparameters, which currently use conventional cytometry settings.

**Figure 8.**
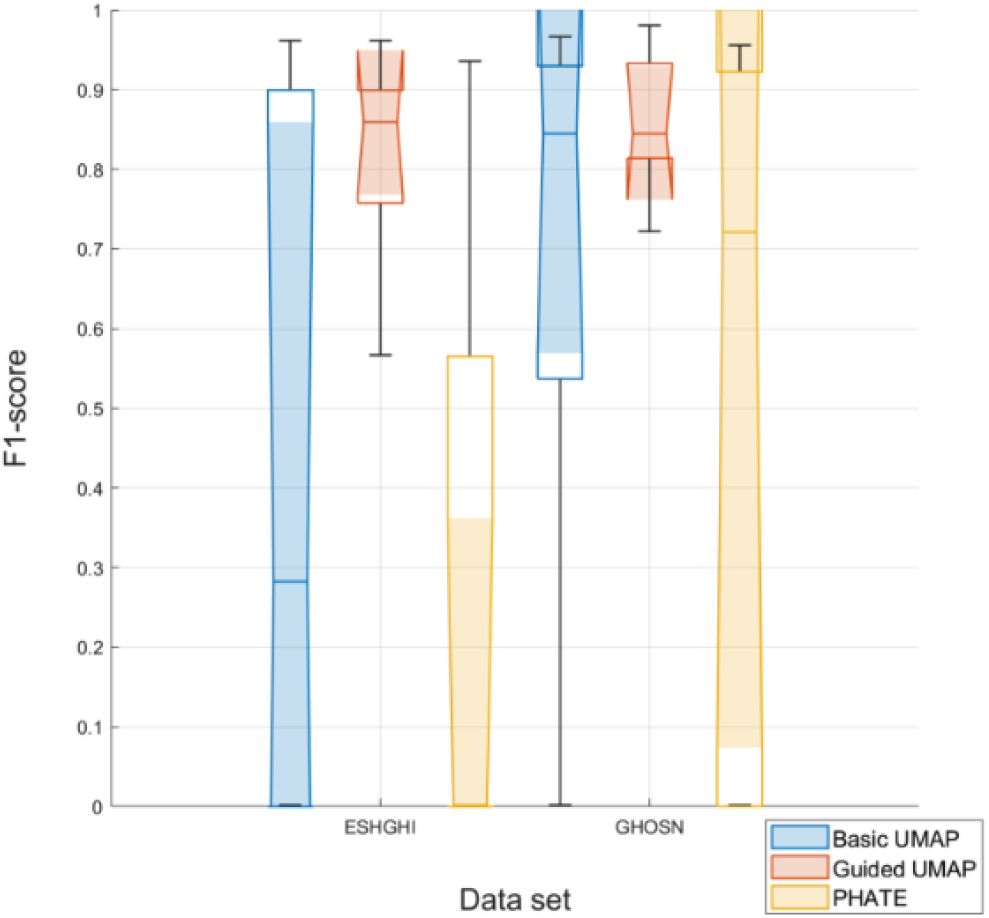
Dimension reduction technique comparison. For both the GHOSN and ESHGHI datasets and each of the three dimension reduction techniques, a box-and-whisker plot shows the median and quartiles of the F1-scores for the dataset’s new populations (those not found in the training dataset), as determined by the original publication’s manual gating. Guided UMAP has the highest median score for both datasets and successfully finds each new population.

## Conclusions

With nine very different published datasets, MLPgater is consistently the best at detecting cell populations, particularly their dense central regions. In six datasets, MLPgater scores remain high for the smallest populations, which often elude even the best automatic gating methods. For example, Eskandari et al.(43) found flowDensity needed “more optimization and potentially incorporating additional tools’’ to confirm some populations containing a small number of cells.

The accuracy of LDA, PhenoGraph and FlowSOM can improve substantially with manual gating additional to that needed to identify live singlets. In contrast, MLPgater performs equivalently well in the presence of dead cells, doublets, off-scale measurements and debris. Hence MLPgater is the easiest to use: simply give it all of a sample’s data and for an assay’s initial setup, also give it each cell’s population label and indicate the FCS measurement parameters (i.e. data columns) used for manual gating (or whatever method is regarded as “ground truth”).

MLPgater spends little more time gating a sample than FlowJo spends copying its manual gates, regardless of sample size or number of gates. Finally, MUDflow not only detects known populations effectively, it also provides a “leg up” advantage for detecting new populations with UMAP’s guided dimensionality reduction feature and DBM clustering. Thus combining these three components creates a complete pipeline for identifying known and new cell populations (both big and small) accurately, quickly and easily in mass and fluorescence cytometry data.

In the future, we will continue to explore ways to improve MUDflow’s discovery of new populations with new datasets and new tunings of the algorithms. We also intend to explore how well MUDflow can handle identifying cell populations in samples with diverse outcomes, as is quite common in the clinical setting.

The Herzenberg Lab provides a software package (named “FlowJoBridge”) capable of running MUDflow free of charge at www.CytoGenie.org. To use either its GUI or API as open source in MATLAB, download it from MathWorks File Exchange(44), unzip it and type “fjb” in the zip file’s root folder. This launches FlowJoBridge, which also accesses this paper’s nine datasets via “demo” FlowJo workspaces.

## Data availability

The raw data files for all datasets used in this paper and the FlowJo Workspace files containing the manual gates used in our testing are available at the following URLs:

**OMIP-044:** Raw data: http://flowrepository.org/id/FR-FCM-ZYC2

Gated: https://storage.googleapis.com/cytogenie.org/Samples/omip44/omip044.wsp

**OMIP-047:** Raw data: http://flowrepository.org/id/FR-FCM-ZYFB

Gated: https://storage.googleapis.com/cytogenie.org/Samples/omipB/omip047_B_cells.wsp

**OMIP-058:** Raw data: http://flowrepository.org/id/FR-FCM-ZYRN

Gated: https://storage.googleapis.com/cytogenie.org/Samples/OMIP-058/OMIP-058.wsp

**OMIP-069:** Raw data: http://flowrepository.org/id/FR-FCM-Z2QV and http://flowrepository.org/id/FR-FCM-Z2QT

Gated: https://storage.googleapis.com/cytogenie.org/Samples/OMIP40Color/OMIP-069_v2.wsp

**OMIP-077:** Raw data: http://flowrepository.org/id/FR-FCM-Z3HC

Gated: https://storage.googleapis.com/cytogenie.org/Samples/OMIP-077/OMIP-077-3WAM.wsp

**ESHGHI:** Raw data: http://flowrepository.org/id/FR-FCM-Z24F

Gated: https://storage.googleapis.com/cytogenie.org/Samples/genentech/Genentech10Merged.wsp

**GHOSN:** Raw data: https://storage.googleapis.com/cytogenie.org/GetDown2/domains/FACS/demo/bCellMacrophag%eDiscovery/all3.zip

Gated: https://storage.googleapis.com/cytogenie.org/GetDown2/domains/FACS/demo/bCellMacrophag eDiscovery/ghosnMacrophagesBCells.wsp

**LEIPOLD:** Raw data: http://flowrepository.org/id/FR-FCM-ZY3Z

Gated: https://storage.googleapis.com/cytogenie.org/Samples/maecker/LEIPOLD.wsp

**PANORAMA:** Raw data: https://storage.googleapis.com/cytogenie.org/Samples/Nikolay/non-Neutrophils.zip Gated: https://storage.googleapis.com/cytogenie.org/Samples/Nikolay/PANORAMA_v3.wsp

## Code availability

The source code for FlowJoBridge can be downloaded from the MathWorks File Exchange at https://www.mathworks.com/matlabcentral/fileexchange/129759-flowjobridge.

## Acknowledgements

We thank Dylan Hinson for support using FlowJo plugins. We thank Sofie Van Gassen for support with analysis and scrutiny of FlowSOM output.

The authors have no funding information to disclose.

## Author contributions

C.M., S.M., J.E., W.M. and L.A.H. contributed to the study conception and design. C.M., S.M. and J.E. were responsible for the software implementation of the algorithms designed in this paper. S.M., W.M. and D.P. contributed to the data curation. C.M., S.M., W.M., D.P., G.W. and L.A.H. participated in the data analysis and interpretation. C.M., S.M. and L.A.H. drafted the article. All authors critically revised the article for important intellectual content.

## Additional information

Supplementary information accompanies this paper. For the present reviewers, please see the submitted PDF file and other files.

### Competing interests

The authors declare no competing interests.

## Notes

### Competing Interest Statement

The authors have declared no competing interest.

