## Supplementary Materials for "MUDflow: Combining Neural Networks, UMAP and DBM Clustering to Identify Cell Populations Accurately, Quickly and Easily in Mass and Fluorescence Cytometry"

### Section S1: Datasets description

To enable an undistracted bioinformatics focus, we evaluate our pipeline using the manual gating and datasets shown in 9 peer-reviewed biology publications. Five of these datasets are collected on conventional cytometers, three on mass cytometers, and one on a spectral cytometer.
The first five datasets listed below are OMIP protocols, which are special peer-reviewed Cytometry Part A publication-type optimized multicolor panels for flow cytometry.
See the "Data availability" section in the paper for access to the original datasets as well as FCS files containing the data with our manual gates.

#### OMIP-044

OMIP-044(1) is a standard 28-color protocol for the human dendritic cell compartment on a conventional cytometer.

1. Manual gating uses 14 markers to define 13 cell populations.
2. LDA and MLPgater test with 3-fold cross-validation over 3 samples. We use 3 of their 6 samples which each contain 2.5 million cells:
    i. samples_AllCells A5807 part1_030.fcs
    ii. samples_FMO1_CD40_033.fcs
    iii. samples_FMO2_CD40_033.fcs

3. Automatic gating operates on the live singlet data starting at the ***CD45+ live*** gate.
4. The first sample gated is samples_AllCells A5807 part1_030.fcs.
5. 14 phenotyping markers define the leaves of the ***CD45+ live*** branch gate**:** CCR7, CD1c, CD3, CD4, CD8, CD11c, CD14, CD16, CD19, CD45RA, CD56, CD123, CD141 and HLA-DR.
6. 13 leaf gates are drawn, following Figure 1 on p. 404 of the publication(1).

### OMIP-047

OMIP-047(2) is a standard 16-color protocol that does dimensional phenotypic characterization of B cells on a conventional cytometer.

1. Manual gating uses 9 markers to define 8 cell populations.
2. LDA and MLPgater test with 4-fold cross-validation over 6 samples (two groups of two samples, two groups of one sample). We use 6 of their 7 samples, omitting 150120_AK156 070309 late chronic_031.fcs.
3. Automatic gating operates on the live singlet data starting at the gate "***B cells"***.
4. The first sample gated is **150416_HC13_037.fcs**.
5. 9 phenotyping markers define the leaves of the **B cells** branch gate: CD10, CD19, CD27, CD38, IgA, IgD, IgG1, IgG3, FSC-A.
6. 8 leaf gates are drawn, following Figure 1 on p. 593 of the publication(2).

### OMIP-058

OMIP-058(3) is a standard 28-color protocol that characterizes NKT, NK, unconventional, and conventional T Cells on a conventional cytometer.

1. Manual gating uses 17 markers to define 35 cell populations.
2. LDA and MLPgater test with 2-fold cross-validation over 2 samples. We use both of their 2 samples, which each contain approximately 1 million cells.
3. Automatic gating operates on the live singlet data starting at the parent gate ***Viable cells***.
4. The first sample gated is "**PBMC_NK and T cell_Donor_112117_D6_D06_015.fcs**".
5. 17 phenotyping markers define the leaves of the ***Viable cells*** branch gate**:** CCR7, CD1d, CD3, CD4, CD8, CD16, CD27, CD28, CD45RA, CD56, CD95, CD161, HLA-DR, TCR Va7_2, TCR Vd1, TCR Vd2, TCR Vg9.
6. 35 leaf gates are drawn, following Figure 1 on p. 948 of the publication(3).

### OMIP-069

OMIP-069(4) is a standard 40-color standard protocol for deep immunophenotyping in human peripheral blood on a spectral cytometer.

1. Manual gating uses 26 markers to define 46 cell populations.
2. LDA and MLPgater test with 3-fold cross-validation over 4 samples (one group of two samples, two groups of one sample). We use all 4 of their samples that each contain approximately 640,000 cells.
3. Automatic gating operates on the live singlet data starting at the parent gate "***CD45+"***.
4. The first sample gated is **MC 3014559.fcs**.
5. **26** phenotyping markers define the leaves of the ***CD45+*** branch gate: CCR7, CD1c, CD2, CD3, CD4, CD8, CD11c, CD14, CD16, CD19, CD20, CD27, CD28, CD38, CD45RA, CD56, CD123, CD127, CD141, CXCR3, CXCR5, HLA-DR, IgD, IgG, IgM, TCRgd,
6. 46 leaf gates are drawn, following Figure 1 on p. 1046 of the publication(4). We separate out 3 gates that are highly overlapping.

### OMIP-077

OMIP-077(5) is a standard 12-color protocol for defining principal human leukocyte populations on a conventional cytometer.

1. Manual gating uses 14 markers to define 14 cell populations
2. LDA and MLPgater test with 4-fold cross-validation over 4 samples. We use all 4 of their samples which contain 7, 3, 1.4 and 2.3 million cells.
3. Automatic gating operates on the live singlet data starting at the parent gate ***Leukocytes***.
4. The first sample gated is **5387 VB IT 080420_Tube_001.fcs**.
5. **14** phenotyping markers define 14 leaves of the ***Leukocytes*** branch gate**:** CD1c, CD3, CD14, CD15, CD16, CD19, CD34, CD38, CD56, CD123, CD141,CD193, HLA-DR, SSC-A.
6. 14 leaf gates are drawn, following Figure 1 on p. 17 of the publication(5).

### ESHGHI

ESHGHI(6) is a study of quantitative differences with manual and t-SNE populations from different sites on a mass cytometer

1. Manual gating uses 27 markers to define 29 cell populations.

2. LDA and MLPgater test with 5-fold cross-validation over 10 samples (five groups of two samples). We use all 10 of their samples.
3. Automatic gating operates on the live singlet data starting at the parent gate "***Residual***".
4. **27** phenotyping markers define 28 leaves of the ***Residual*** branch gate**:** CCR7, CD1c, CD3, CD4, CD8, CD11b, CD11c, CD14, CD16, CD19, CD20, CD25, CD27, CD38, CD45, CD45RA, CD49d_alpha4, CD56, CD66, CD123, CD161, FcEri, Foxp3, gamma_delta_TCR, HLADR, IgD, Va7.2.
5. 28 leaf gates are drawn, following Figure 1 on p. 4 of the publication(6).

### GHOSN

GHOSN(7) is a study of macrophages and B cells in the peritoneal cavity of 3 strains of mice each stained with a slightly different reagent cocktail on a <u>conventional cytometer</u>.

1. Manual gating uses 9 markers to define 13 cell populations.
2. LDA and MLPgater test with 2-fold cross-validation over 3 samples (one group of two samples, one group of one sample). We use 3 of their 4 samples which each contain 200,000 cells:
   i. "all_3-1.fcs" (C57 strain)
   ii. "all_3-2.fcs" (BALB/c strain with fluorescence-minus-one for Gr-1: APC)
   iii. "all_3-3.fcs" (BALB/c strain)
3. Automatic gating operates on the live singlet data starting at the parent gate "***Live singlets***".
4. The base sample gated is **all_3-3.fcs**.
5. **9** phenotyping markers define 13 leaves of the ***Live singlets*** branch gate**:** CD5, CD19, CD11c, CD11b, F4/80, IgD, IgM, FSC-A, SSC-A.
6. 13 leaf gates are drawn, following Figure 1 on p. 2569 of the publication(7).

### LEIPOLD

LEIPOLD(8) is a study comparing CyTOF assays from 6 sites on a <u>mass cytometer</u>.

1. Manual gating uses 15 markers to define 18 cell populations.
2. LDA and MLPgater test with 4-fold cross-validation over 6 samples (two groups of two samples, two groups of one sample). We use all 6 of their samples, which range in size from 29,713 to 112,843 cells.
3. Automatic gating operates on the live singlet data starting at the parent gate "***All cells***".

4. The base sample gated is **Center_1_A_1_MATLAB_LiveInactSing_Linneg_CD45pos.fcs**.
5. **15** phenotyping markers define 18 leaves of the ***All cells*** branch gate**:** CD3, CD4, CD8A, CD11c, CD14, CD16, CD19, CD20, CD27, CD33, CD38, CD45RA, CD56, CD123, and HLADR.
6. 18 leaf gates are drawn, following Supplementary Fig. S6 of the publication(8).

#### PANORAMA

PANORAMA(9) is an often-cited dataset collected on the mass cytometer. It is frequently used for testing gating methods. The publication that introduces the X-Shift classifier, which explores this dataset.

1. Manual gating uses 25 markers to define 24 cell populations.
2. LDA and MLPgater test with 5-fold cross-validation over 10 samples (five groups of two samples). We use all 10 of their samples, which range in size from 75,513 to 95,087 cells.
3. Automatic gating operates on the live singlet data starting at the parent gate ***non-Neutrophils***.
4. The base sample gated is **BM2_cct_normalized_01_non-Neutrophils.fcs**.
5. **25** phenotyping markers define 24 leaves of the ***non-Neutrophils*** branch gate**:** 120g8, B220, CD3, CD4, CD8, CD11b, CD11c, CD19, CD34, CD43, CD49b, CD64, CD115, CD138, DNA2, F480, FceR1a, IgM, Ly6C, Ly6G, MHCII, NKp46, SiglecF, TCRb, TCRgd.
6. 24 leaf gates are drawn, following Supplementary Fig. 5 of the publication(9).

### Section S2: FlowSOM hyperparameters

Through private correspondence with one of the authors of the original FlowSOM publication(10), Sofie Van Gassen, we learned that optimal results cannot be expected with the guidance given by BD's FlowJo support team and the flowLearn publication(11) when they compare with FlowSOM. Setting "meta clusters" to the number of expected populations plus one and keeping the "grid size" at 10 is not optimal. The "meta clusters" and "grid size" hyperparameters given to FlowSOM do need testing and tuning per assay, but a better yet rough starting point for "meta clusters" is 150% of the expected populations rounded down. For "grid size", a better starting point is the square root of 5 times the meta clusters rounded down. The minimum grid size is 10. We leave all other hyperparameters in the FlowJo 10.10 plugin configuration screen unaltered from their default values.

### Section S3: Pre-filtering datasets

The data preparation pattern practiced by Abdelaal et al.(12) involves first removing all cells from all datasets that do not belong to any population defined by manual gating. We confirmed this by inspecting both the code (available at https://github.com/tabdelaal/CyTOF-Linear-Classifier) as well as the datasets referenced in the publication.

To compare directly with their processing, we performed a round of testing both MLPgater and LDA by first preparing each sample in all of our 9 datasets by removing the unlabeled cells for both training and predicting. We generalized Abdelaal et al.'s multiple dataset-specific scripts into one generic one and then plugged in our MLPgater alongside their LDA. As a quality control check, we reran LDA with these modified scripts on the 6 CyTOF datasets described in their publication in order to reproduce their published results and thus confirm that our changes did not introduce problems. A summary of the testing results is shown in Tables S1 and S2.

The testing results indicate that automatically classified populations match better to manually gated populations when unclassified cells are excluded from the computation.

| ***Classification Method*** | ***Data Included in Computation*** | *F1-Score* | | *CS* | |
|---|---|---|---|---|---|
| | | *Median* | *Mean* | *Median* | *Mean* |
| MLPgater | Classified only | 0.99 | 0.95 | 0.99 | 0.94 |
| MLPgater | All | 0.98 | 0.91 | 0.98 | 0.89 |
| LDA | Classified only | 0.88 | 0.81 | 0.83 | 0.76 |
| LDA | All | 0.80 | 0.72 | 0.74 | 0.66 |

**Table S1. Summary of effect of removing manually unclassified cells from datasets.**

| Dataset | MLPgater | | | | LDA | | | |
|---|---|---|---|---|---|---|---|---|
| | Classified only | | Classified+unclassified | | Classified only | | Classified+unclassified | |
| | *F1-Score* | *Central similarity* | *F1-Score* | *Central similarity* | *F1-Score* | *Central similarity* | *F1-Score* | *Central similarity* |
| OMIP-044 | 1<br>*0.99* | 1<br>*l* | 1<br>0.99 | 1<br>0.99 | 0.93<br>*0.85* | 0.90<br>*0.81* | 0.91<br>*0.84* | 0.89<br>**0.80** |
| OMIP-047 | 1<br>*1* | 1<br>*1* | 0.98<br>0.97 | 0.99<br>0.98 | 0.96<br>*0.96* | 0.95<br>*0.96* | **0.88**<br>0.85 | 0.87<br>0.83 |
| OMIP-058 | 0.99<br>*0.97* | 0.99<br>*0.97* | 0.98<br>0.97 | 0.98<br>0.96 | 0.84<br>***0.76*** | 0.81<br>*0.70* | **0.75**<br>0.70 | 0.62<br>0.61 |
| OMIP-069 | 0.98<br>*0.93* | 0.97<br>**0.90** | **0.87**<br>***0.76*** | 0.81<br>*0.70* | 0.83<br>***0.76*** | 0.73<br>**0.68** | **0.58**<br>0.56 | 0.45<br>0.46 |
| OMIP-077 | 0.97<br>*0.89* | 0.97<br>*0.85* | **0.93**<br>***0.84*** | 0.93<br>*0.79* | 0.72<br>*0.62* | 0.62<br>***0.57*** | 0.70<br>*0.57* | 0.62<br>*0.53* |
| ESHGHI | 1<br>*0.99* | 1<br>*0.99* | 0.98<br>0.97 | 0.99<br>0.97 | 0.90<br>0.86 | 0.86<br>***0.81*** | 0.87<br>**0.82** | 0.82<br>**0.77** |
| GHOSN | 0.99<br>*0.94* | 0.99<br>***0.92*** | 0.97<br>0.95 | 0.98<br>0.95 | 0.92<br>0.89 | 0.93<br>***0.85*** | **0.84**<br>***0.82*** | 0.84<br>*0.78* |
| LEIPOLD | 0.99<br>*0.96* | 0.99<br>*0.95* | 0.98<br>0.94 | 0.98<br>0.92 | 0.86<br>***0.78*** | 0.80<br>***0.73*** | **0.80**<br>***0.69*** | 0.74<br>*0.64* |
| PANORAMA | 0.97<br>*0.88* | 0.97<br>*0.86* | **0.93**<br>***0.81*** | 0.89<br>*0.76* | 0.88<br>*0.79* | 0.83<br>*0.74* | **0.65**<br>0.62 | 0.51<br>0.53 |

**Table S2. Effect of removing manually unclassified cells from datasets.**

All datasets except OMIP-077 LDA's predictive accuracy improves significantly more than MLPgater's predictive accuracy improves. Moreover, there are only 3 of 9 (OMIP-047, OMIP-058 and PANORAMA) datasets where MLPgater's F1-score improves by more than 0.01. In these 3 datasets, we also observe significantly lower central similarity scores telling us that MLPgater sees the dense central regions of phenotypes in N-dimensional space more clearly for these assays when unclassified cells are removed.

Pursuing improvements in this manner however is non-ideal because while removing unclassified cells may be fair and easy for training, predicting on such data requires a prior pregating step that makes it difficult to deprecate manual gating. Therefore, in this paper, we draw our conclusions on test results where unclassified cells are included.

### Section S4: Many-to-many matching

Our QFMatch(13) software provides 4 options for merging

1. Merging subsets in both the training set and in the test set, which allows for *many-to-many matching.*
2. Merging subsets only in the test set which allows for *one-to-many matching.*
3. Merging subsets only in the training set which allows for *many-to-one matching.*
4. No merging.

We performed a round of testing both FlowSOM and PhenoGraph while using many-to-many matching in step 3c of “Testing procedures” in the paper. The testing results in Tables S3 and S4 indicate that unsupervised automatically classified populations match better to manually gated populations when many-to-many matching is used.

| **Performance metric** / **Classifier** | *F1-score* | | *Central Similarity* | |
|---|---|---|---|---|
| | *Median* | *Mean* | *Median* | *Mean* |
| FlowSOM | | | | |
| Many-to-many | 0.87 | 0.74 | 0.86 | 0.70 |
| One-to-many | 0.61 | 0.48 | 0.48 | 0.44 |
| PhenoGraph | | | | |
| Many-to-many | 0.87 | 0.72 | 0.84 | 0.66 |
| One-to-many | 0.76 | 0.60 | 0.69 | 0.55 |

**Table S3. Summary of effect of many-to-many matching for unsupervised methods.**

| Dataset | FlowSOM | | | | PhenoGraph | | | |
|---|---|---|---|---|---|---|---|---|
| | Many-to-many | | One-to-many | | Many-to-many | | One-to-many | |
| | F1-score | Central similarity | F1-score | Central similarity | F1-score | Central similarity | F1-score | Central similarity |
| OMIP-044 | .97<br>*.88* | .98<br>***.86*** | 0.61<br>*0.49* | 0.51<br>*0.46* | .92<br>***.81*** | .92<br>***.78*** | 0.91<br>*0.81* | 0.92<br>*0.78* |
| OMIP-047 | .87<br>*.86* | .83<br>*.81* | 0.76<br>*0.60* | 0.67<br>*0.55* | .90<br>*.83* | .87<br>***.77*** | 0.89<br>*0.83* | 0.87<br>*0.77* |
| OMIP-058 | .80<br>***.60*** | .73<br>***.55*** | 0.44<br>*0.42* | 0.31<br>*0.38* | .82<br>***.63*** | .77<br>***.58*** | 0.48<br>*0.44* | 0.77<br>*0.58* |
| OMIP-069 | .51<br>*.48* | .37<br>*.40* | 0.31<br>*0.35* | 0.20<br>*0.29* | .55<br>*.52* | .41<br>*.43* | 0.28<br>*0.34* | 0.15<br>*0.29* |
| OMIP-077 | .96<br>*.92* | .95<br>*.92* | 0.65<br>*0.52* | 0.55<br>*0.50* | .96<br>*.86* | .96<br>*.86* | 0.91<br>*0.66* | 0.92<br>*0.65* |
| ESHGHI | .94<br>*.84* | .95<br>***.82*** | 0.60<br>*0.49* | 0.48<br>*0.46* | .88<br>***.74*** | .84<br>***.71*** | 0.76<br>*0.61* | 0.69<br>*0.57* |
| GHOSN | .93<br>*.83* | .92<br>***.82*** | 0.81<br>*0.60* | 0.74<br>*0.59* | .87<br>*.79* | .89<br>***.74*** | 0.81<br>*0.71* | 0.74<br>*0.65* |
| LEIPOLD | .87<br>***.72*** | .86<br>***.70*** | 0.21<br>*0.40* | 0.11<br>*0.36* | .84<br>***.68*** | .80<br>***.64*** | 0.70<br>*0.60* | 0.61<br>*0.54* |
| PANORAMA | .56<br>*.53* | .41<br>*.44* | 0.55<br>*0.45* | 0.40<br>*0.37* | .56<br>*.59* | .37<br>*.48* | 0.32<br>*0.38* | 0.19<br>*0.31* |

**Table S4. Effect of many-to-many matching for unsupervised methods.**

Because we treat scientist-reviewed manually gated populations as ground truth, we do not allow manual gates to be merged in the matching process in our normal testing procedure.

### Section S5: Small population issues

Automatic gating methods often miss smaller populations important to clinical research(14). In our testing, however, MLPgater's F1-score is at least 0.70 in 6 datasets. In the other 3 datasets,

11 populations have MLPgater F1-scores < 0.50: 1 in OMIP-077 (0.44), 4 in PANORAMA and 6 in OMIP-069.

Table S5 identifies the population with the minimum F1 score for each classifier in each dataset.

| *Minimum F1-score, population name and population % frequency* | | | | |
|---|---|---|---|---|
| **Classifier** / **Dataset** | **MLPgater** | **LDA** | **PhenoGraph** | **FlowSOM** |
| OMIP-044 | 0.901<br>*CD141+ DCs* 0.04% | .373<br>*DCs*.1.4% | .197<br>*DCs*.1.4% | 0<br>*not naive CD4+*.18%<br>***and 4 others*** |
| OMIP-047 | 0.881<br>*Transitional* 2.2% | .613<br>*IgA, FSC-A IgA, IgG1, IgG3* 7.3% | .549<br>*PB* 0.38% | 0<br>*Transitional* 2.2%<br>*IgA, FSC-A IgA+* 5.5% |
| OMIP-058 | 0.902<br>*Terminal NK*.1% | 0.167<br>CD4+ TSCM T cells CD27+ CD28+ 1% | 0<br>*CD4+ TEMRA T Cells CD27- CD28+* 0.03%<br>***and 12 others*** | 0<br>*CD4+ TEMRA T Cells CD27- CD28+*.03%<br>***and 12 others*** |
| OMIP-069 | 0<br>*CD1c- CD141+ DCs* **0.01%** | 0.17<br>*CD16- CD14- monocytes* .0.30% | 0<br>*B-1 IgM+* 0.21%<br>***and 13 others*** | 0<br>*B-1 IgM+* 0.21%<br>***and 8 others*** |
| OMIP-077 | 0.442<br>*pDCs* 0.01% | 0.022<br>*CD141+ mDCs* 0.04% | 0<br>*CD141+ mDCs* 0.41%<br>***and 2 others*** | 0<br>*CD141+ mDCs* 0.41%<br>***and 4 others*** |
| ESHGHI | 0.895<br>*CD123-* 0.17% | 0.430<br>*Lineage Negative* 0.57% | 0<br>*CD123-* 0.17%<br>***and 3 others*** | 0<br>*CD123-* 0.17%<br>***and 7 others*** |
| GHOSN | 0.854<br>*Neutrophils* 0.36% | 0.394<br>*Neutrophils* 0.36% | 0<br>*CD11b+ dendritic* 1.3% | 0<br>*ckit+ mast cells* 1.8%<br>***and 3 others*** |
| LEIPOLD | 0.697<br>*CD16+ monocytes* 0.04% | 0.055<br>*CD16+ monocytes* 0.04% | 0<br>*CD8 T cells CD45RA- CD27-* 2.8% | 0<br>*CD16+ monocytes* 0.04%<br>***and 4 others*** |
| PANORAMA | 0<br>*HSC* 0.02% | 0<br>*HSC* 0.02% | 0<br>*HSC* 0.02%<br>***and 6 others*** | 0<br>*gd T cells* 0.21%<br>***and 3 others*** |

**Table S5. Minimum-scoring population for each classifier and dataset.**

MLPgater is the only classifier having datasets (5 of 9) where the worst F1-score still exceeds 0.85: OMIP-044, OMIP-047, OMIP-058 and ESHGHI. Moreover, for 197 populations overall MLPgaters F1-Score drops below 0.50 11 times.

In Figure S1, the F1-Score by Size X/Y plots of all classifiers compared side by side illustrate the strength of MLPgater with small populations. Even in the two worst cases (PANORAMA and OMIP-069), the plots indicate that MLPgater is much less affected by population size than are LDA, FlowSOM and PhenoGraph. Even in the two worst cases (PANORAMA and OMIP-069), the plots below indicate that overall MLPgater is much less affected by population size than are LDA, FlowSOM and PhenoGraph.

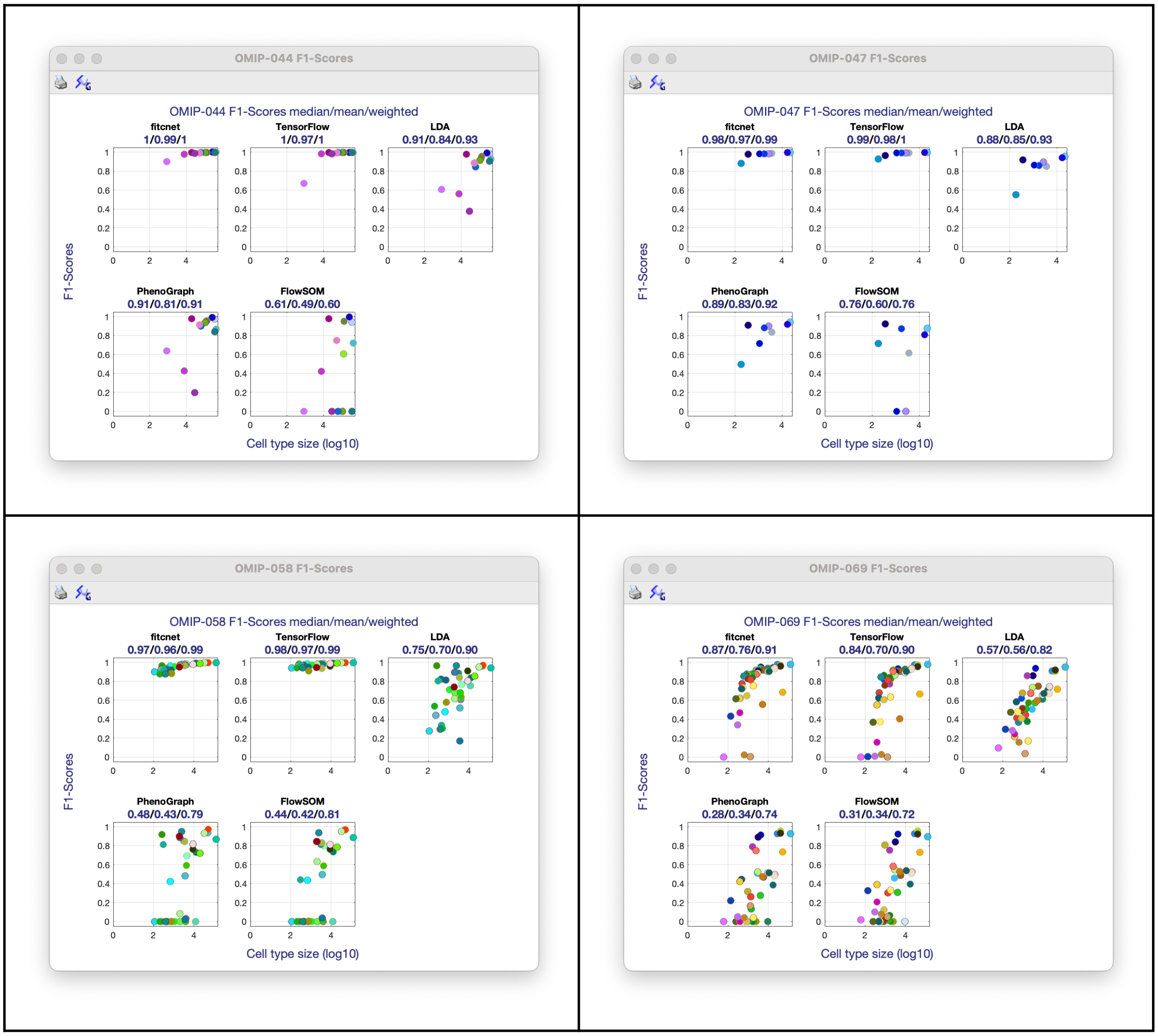

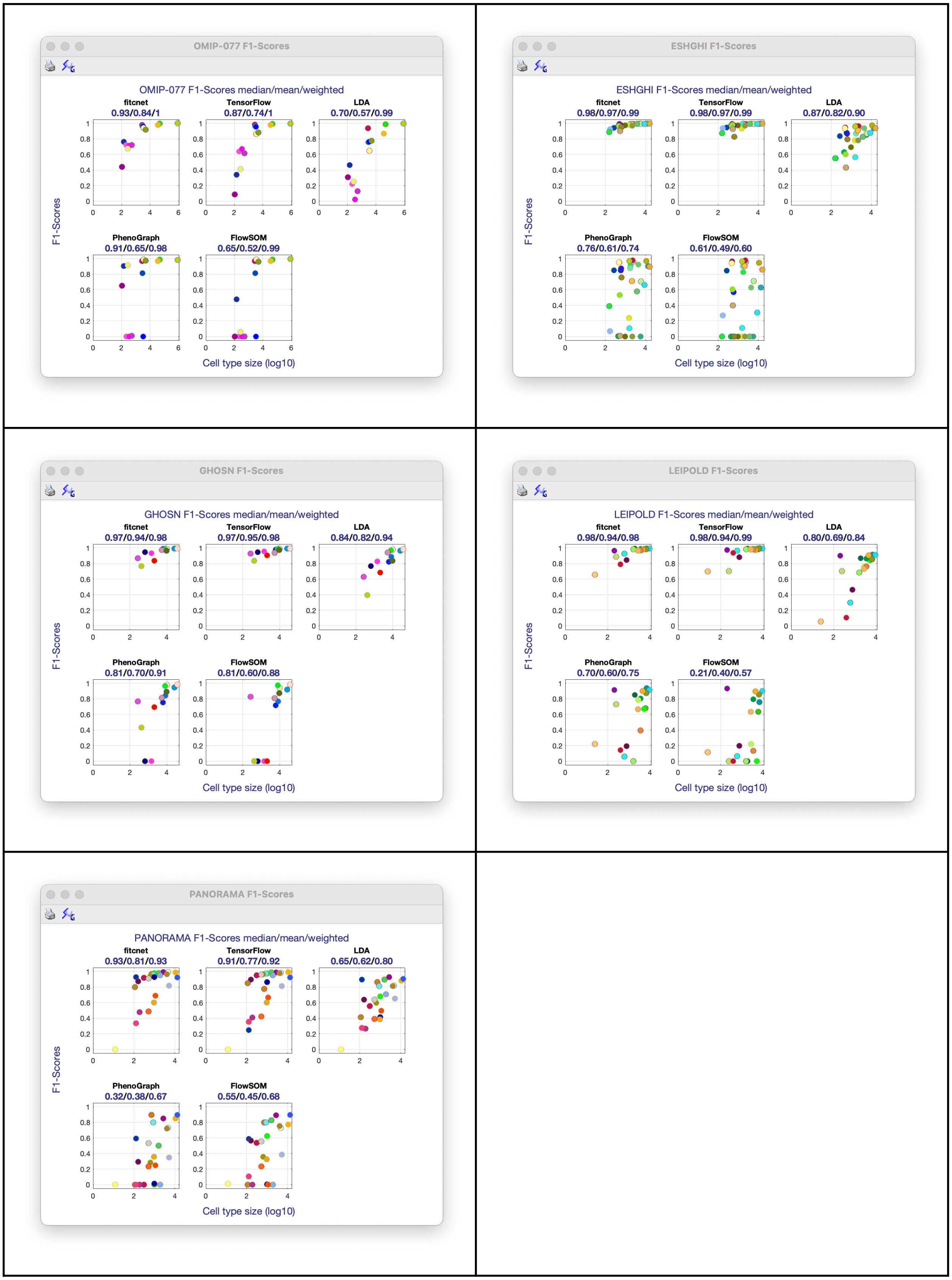

**Figure S1. F1-score vs. Population size for each classifier and dataset.**

### Section S6: Low scores in OMIP-069

Our population level results in this dataset stand out because

1. This is where MLPgater's scores lowest.
2. Nevertheless, the gap between MLPgater scores and those of LDA, FlowSOM and PhenoGraph is the biggest

Concerning item 1 above, the matches that are atypically poor for MLPgater occur on cell populations whose manual gate sequence contains drawings that are also atypical by not enclosing the density features that probability coloring and contours highlight in the X/Y plots plots reflecting prior biology knowledge beyond the data's structure. If we ignore such regions (e.g., monocytes) and instead focus on a region like that containing the 19 leaf gates under the parent gates "T cells" and "NKT-like cells", then we find that MLPgater achieves more typical high scores, as shown in Table S6.

| Color | Cell type | Frequency % | F1-Score | | Central similarity | |
|---|---|---|---|---|---|---|
| | | | fitcnet | LDA | fitcnet | LDA |
| | Naive CD4+ | 46% | 0.995 | 0.979 | 0.999 | 0.997 |
| | Naive CD8+ | 15% | 0.994 | 0.945 | 0.999 | 0.978 |
| | Early like effector CD4+ | 3.4% | 0.980 | 0.676 | 0.981 | 0.563 |
| | Early Effector CD4+ | 6.4% | 0.979 | 0.796 | 0.979 | 0.725 |
| | Central Memory CD4+ CXCR5-, CXCR3 | 7.1% | 0.964 | 0.761 | 0.962 | 0.682 |
| | Central Memory CD4+ CXCR5-, CXCR3- | 7.6% | 0.954 | 0.785 | 0.939 | 0.709 |
| | Early Effector CD8+ | 4% | 0.945 | 0.811 | 0.920 | 0.736 |
| | Intermediate Effector CD8+ | 0.70% | 0.943 | 0.719 | 0.924 | 0.620 |
| | Central Memory CD8+ | 0.53% | 0.943 | 0.638 | 0.915 | 0.485 |
| | Terminal Effector CD4+ | 0.22% | 0.933 | 0.412 | 0.918 | 0.281 |
| | CD4- CD8- | 1.1% | 0.930 | 0.710 | 0.885 | 0.560 |
| | Terminal Effector | 0.31% | 0.909 | 0.571 | 0.866 | 0.458 |
| | Early like effector CD8+ | 0.59% | 0.902 | 0.616 | 0.853 | 0.493 |
| | Terminal Effector CD45RA+ CD8+ | 1.5% | 0.885 | 0.677 | 0.814 | 0.572 |
| | Central Memory CD4+ CXCR5, CXCR3 | 3.4% | 0.863 | 0.724 | 0.809 | 0.617 |
| | CD4+ CD8+ T | 0.17% | 0.844 | 0.700 | 0.797 | 0.562 |
| | CD2- NKT cells | 0.09% | 0.154 | 0.254 | 0.097 | 0.158 |
| | CD2+ CD8+ NKT | 2.1% | 0.053 | 0.120 | 0.025 | 0.067 |
| | CD2+ CD8- | 0.34% | 0.029 | 0.050 | 0.015 | 0.030 |
| | | **Median** | **0.933** | **0.700** | **0.915** | **0.563** |
| | | **Mean** | **0.800** | **0.629** | **0.774** | **0.542** |

**Table S6. Improved performance metrics for MLPgater and LDA after classifying over a smaller region of cells.**

Concerning item 2, MLPgater's significantly greater accuracy also occurs with the PANORAMA and OMIP-058 datasets. We suspect there must be more tuning with each of these classifiers' configuration parameters than those settings which FlowJo's support team recommended with PhenoGraph and FlowSOM plugins or that which the LDA publication's supplementary code illustrated. The OMIP-069 publication itself discusses the use of FlowSOM on their data. There may be better tunings of FlowSOM, LDA or PhenoGraph that are more accurate for this spectral cytometry dataset as well as for the others, but this means that they do not meet one of our bioinformatics goals of being easy to use for the biologist as well as accurate and quick. Exploring this further for OMIP-069, we pointed FlowSOM at the same data region upon which we narrowed MLPgater's attention for the same reasons. This region contains 19 populations, so we set FlowSOM to find 20 meta clusters to allow 1 for background and then ran it producing the results shown in Table S7 and discussed below. The median/mean of the F1-score were 0.405/0.410 which improves upon the 0.306/0.346 obtained for running FlowSOM on all 46 populations. But this does not inspire trust in the capability of FlowSOM.

If QFMatch is run to find the best many-to-many F1-score between FlowSOM gated populations and manually gated populations, however, the median/mean goes from 0.31/0.35 to 0.511 /0.481. QFMatch finds 7 of 30 matches where merging multiple manual or FlowSOM gated populations obtains a higher F1-score. The 10th, 18th, 21st and 26th ranked matches merge multiple manually gated populations to match one FlowSOM gated population. The 2nd, 5th and 12th ranked matches merge multiple FlowSOM gated populations to match one manually gated population.

We run FlowSOM with 44 meta clusters to find 43 cell populations (the one extra is for background). But FlowSOM only finds 42 clusters. So we tried running FlowSOM on only the T cells and NKT-like cells which require 9 phenotyping markers that classify 19 leaf gates starting at the CD3+ TCRgd- branch gate: CCR7, CD2, CD4, CD8, CD27, CD28, CD45RA, CXCR3, CXCR5. We ran FlowSOM on the common parent gate to these populations specifying 20 meta clusters to allow 1 for background. The median/mean of the F1-score shown below are 0.405/0.410. This improves upon the 0.306/0.346 obtained for running FlowSOM on all 46 populations. The same populations found by FlowSOM when starting at the "CD45+" branch closest to all 43 cell populations get lower scores than when FlowSOM starts at the lower CD3+ TCRgd- branch: for example, Naive CD8+ gets 0.90 F1-score on the CD45+ branch run and 0.98 on the run for the branch with only NKT and T leaves.

| Color | Cell type | Frequency % | F1-Score | Central similarity |
|---|---|---|---|---|
| | | | FlowSOM | FlowSOM |
| • | Naive CD8+ | 18% | 0.979 | 0.996 |
| • | Naive CD4+ | 39% | 0.950 | 0.959 |
| • | CD4- CD8- | 2% | 0.776 | 0.651 |
| • | Central Memory CD4+ CXCR5-, CXCR3 | 6.3% | 0.687 | 0.533 |
| • | Early Effector CD8+ | 4.1% | 0.661 | 0.524 |
| • | CD4+ CD8+ T | 0.13% | 0.656 | 0.499 |
| • | Early like effector CD4+ | 2.4% | 0.526 | 0.385 |
| • | Central Memory CD4+ CXCR5-, CXCR3- | 8.8% | 0.515 | 0.354 |
| • | Early Effector CD4+ | 5.4% | 0.489 | 0.328 |
| • | Intermediate Effector CD8+ | 0.66% | 0.405 | 0.277 |
| • | Terminal Effector CD45RA+ CD8+ | 0.61% | 0.401 | 0.273 |
| • | Early like effector CD8+ | 0.77% | 0.393 | 0.253 |
| • | CD2+ CD8+ NKT | 2.3% | 0.280 | 0.187 |
| • | Central Memory CD8+ | 0.50% | 0.071 | 0.033 |
| • | Central Memory CD4+ CXCR5, CXCR3 | 8.8% | 0.003 | 0.001 |
| • | CD2+ CD8- | 0.56% | N/A | N/A |
| • | Terminal Effector | 0.30% | 0 | 0 |
| • | CD2- NKT cells | 0.13% | N/A | N/A |
| • | Terminal Effector CD4+ | 0.01% | N/A | N/A |
| | | **Median** | **0.405** | **0.277** |
| | | **Mean** | **0.410** | **0.329** |

**Table S7. Improved performance metrics for FlowSOM after classifying over a smaller region of cells.**

Running PhenoGraph on the more narrow region of the gating tree at the ***CD3+ TCRgd-*** branch also improves the F1-score median/mean from 0.275/0.338 to 0.414/0.375. Individual populations like CD4- CD8- improves from 0.52 to 0.76. See Table S8.

| Color | Cell type | Frequency % | F1-Score | Central similarity |
|---|---|---|---|---|
| | | | PhenoGraph | PhenoGraph |
| | Naive CD8+ | 18% | 0.97 | 0.99 |
| | Naive CD4+ | 39% | 0.92 | 0.92 |
| | CD4- CD8- | 2% | 0.76 | 0.65 |
| | Central Memory CD4+ CXCR5, CXCR3 | 8.8% | 0.68 | 0.56 |
| | Early like effector CD4+ | 2.4% | 0.61 | 0.46 |
| | Early Effector CD8+ | 4.1% | 0.56 | 0.43 |
| | Central Memory CD4+ CXCR5-, CXCR3 | 6.3% | 0.52 | 0.39 |
| | CD4+ CD8+ T | 0.13% | 0.47 | 0.28 |
| | CD2+ CD8+ NKT | 2.3% | 0.44 | 0.31 |
| | CD2- NKT cells | 0.13% | 0.41 | 0.30 |
| | Early Effector CD4+ | 5.4% | 0.40 | 0.26 |
| | Terminal Effector CD45RA+ CD8+ | 0.61% | 0.26 | 0.16 |
| | Central Memory CD4+ CXCR5-, CXCR3- | 8.8% | 0.09 | 0.05 |
| | Intermediate Effector CD8+ | 0.66% | 0.01 | |
| | Central Memory CD8+ | 0.50% | 0.01 | 0.01 |
| | CD2+ CD8- | 0.56% | 0.01 | 0.01 |
| | Early like effector CD8+ | 0.77% | | |
| | Terminal Effector | 0.30% | | |
| | Terminal Effector CD4+ | 0.01% | | |
| 19 cell populations | | Median | 0.414 | 0.283 |
| | | Mean | 0.375 | 0.303 |

**Table S8. Improved performance metrics for PhenoGraph after classifying over a smaller region of cells.**

### Section S7: Running MLPgater on doublets, debris, and dead cells

On the GHOSN dataset(7), MLPgater runs as effectively on the whole sample as it does starting at the live singlets branch gate. In Figure S2, we compare using the entire raw sample versus only the live singlets while training MLPgater on the BALB/c mouse strain sample and then identifying populations in the C57 mouse strain (all_3-2.fcs).

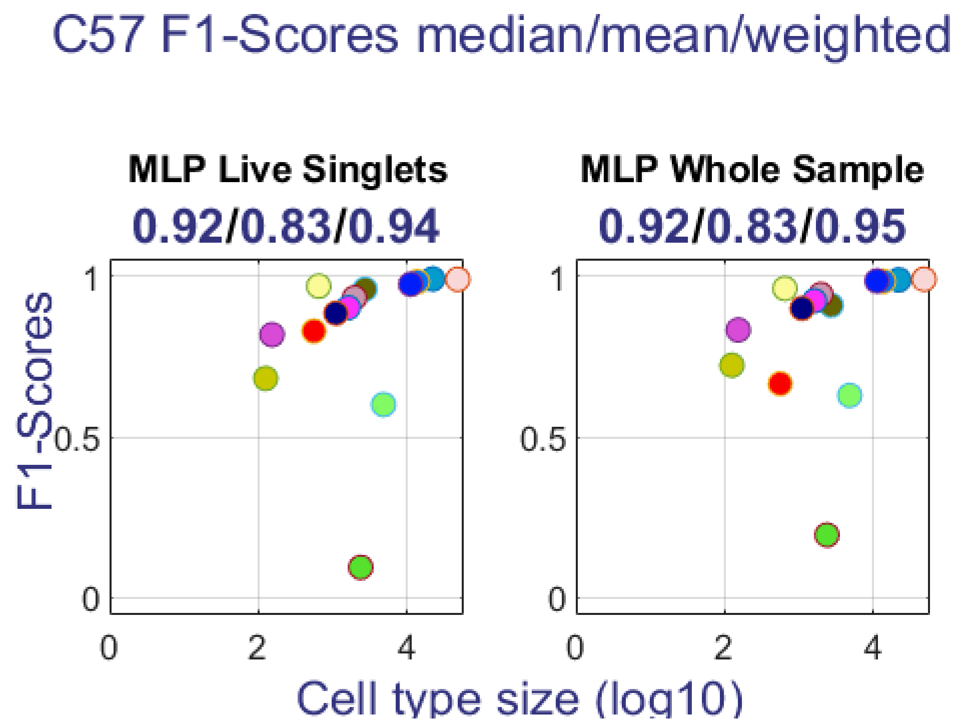

**Figure S2. F1-scores of populations identified by MLPgater when using the whole sample vs. live singlets.**

Guiding UMAP with predictions on the whole C57 sample produces the plot in Figure S3.

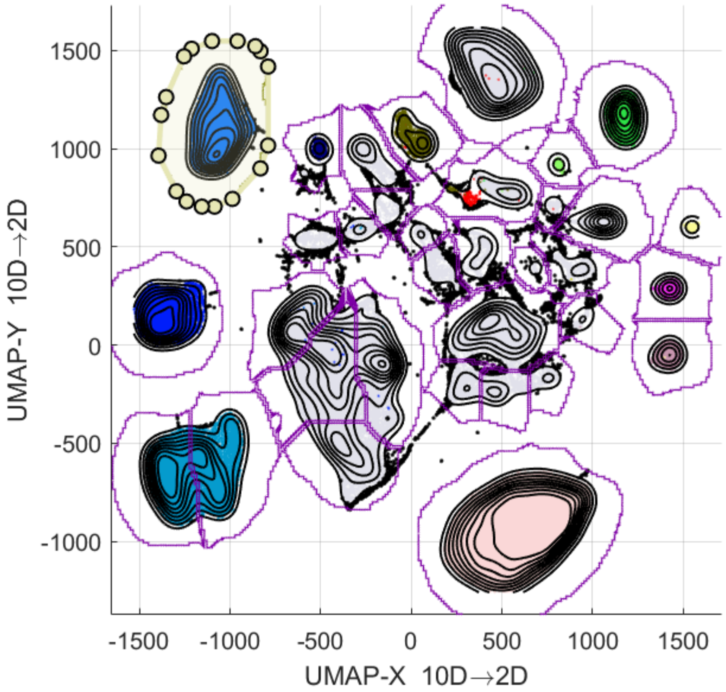

**Figure S3. UMAP plot of C57 mouse sample guided by MLPgater classification over the entire sample.**

Highlighting manual gates for doublets, dead cells and debris locates most of them in the uncolored clusters, as shown in Figure S3.

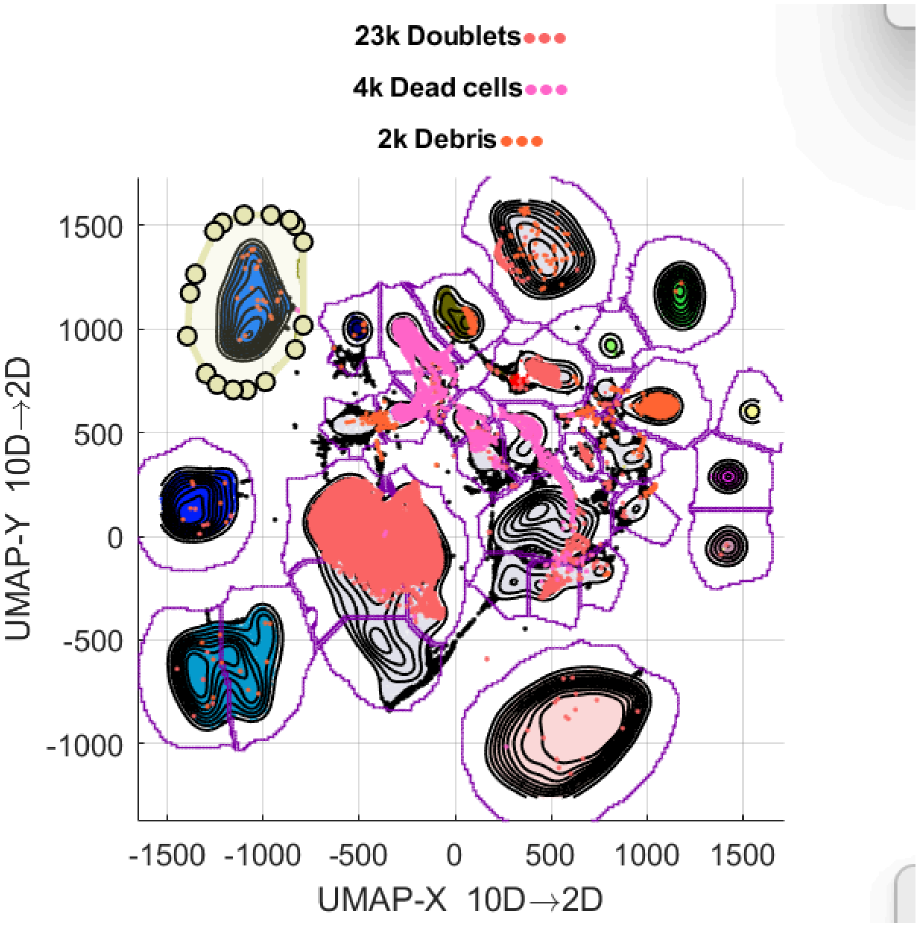

**Figure S3. Doublets, dead cells, and debris in the UMAP plot.**

### Section S8: Basic UMAP plots

Like other dimension reduction techniques, UMAP is primarily designed and used as a visualization tool(15,16). In flow informatics, it can allow researchers to use pattern recognition in order to discover biological insights(17). It is therefore natural to wonder if “basic UMAP” (see Results for an explanation of the difference between “basic UMAP” and “guided UMAP”) can be used directly as a visual tool for identifying new cell populations in new biological samples.

See Figure S4 for an example of two basic UMAP plots of compatible samples from two different strains of laboratory mice. The plot on the right contains biologically novel cell populations corresponding to lymphocytes. Since we are assuming no prior knowledge of cell types, we use colors to signify data density, rather than population identity.

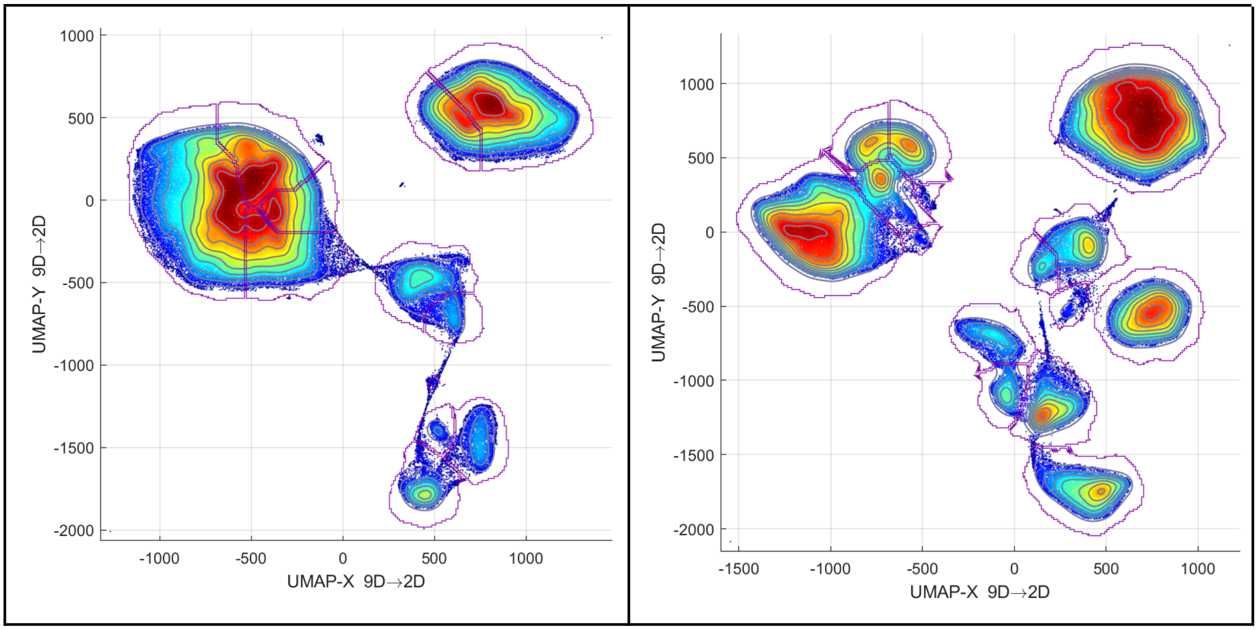

**Figure S4. Basic UMAP plots of cells from a peritoneal cavity of a RAG mouse (left) and BALB/c mouse (right)**

From Figure S4, we can see that although there are some visual similarities between the two plots, there is no direct correspondence between their clusters. For example, through direct inspection of the underlying datasets, we find that the cluster in the top-left of the right plot corresponds to a biologically novel population (B cells), and does not match the cluster in the top-left of the left plot (macrophages).

In general, for two different UMAP dimension reductions $g_1$ on the dataset $B_1$ and $g_2$ on $B_2$, there is little reason to believe that $g_1$ and $g_2$ will preserve global and local structure from high-dimensional space, This is a consequence of the fact that there is no "universal embedding" $g : \mathbb{R}^N \to \mathbb{R}^2$ from which all other UMAP embeddings derive; each UMAP embedding is calculated based on the assumption that the original finite dataset lies on a customized high-dimensional manifold(15).

It is still possible to identify new populations from visual inspection by utilizing additional steps (e.g. viewing "heat maps" of clusters in high-dimensional space to create a correspondence between clusters of different plots). However, such methods would likely be time-consuming and so would require automation to reach our goal of speed.

We investigate one approach for automating discovery of new populations in basic UMAP plots in Results, where we integrate it into the MUDflow pipeline.

### Section S9: Results for populations

#### OMIP-044

| Population | Freq-uency | MLP F1-score | MLP CS | LDA F1-score | LDA CS | PhenoGraph F1-score | PhenoGraph CS | FlowSOM F1-score | FlowSOM CS |
|---|---|---|---|---|---|---|---|---|---|
| CD14+ | 19% | 1.00 | 1.00 | 0.99 | 1.00 | 0.97 | 0.99 | 0.99 | 1.00 |
| CD19+ | 13% | 1.00 | 1.00 | 0.99 | 1.00 | 0.99 | 1.00 | 0.99 | 1.00 |
| naive CD4+ | 21% | 1.00 | 1.00 | 0.91 | 0.87 | 0.87 | 0.79 | 0.72 | 0.57 |
| [not naive CD4+] | 19% | 1.00 | 1.00 | 0.90 | 0.85 | 0.84 | 0.76 | | |
| naive CD8+ | 6.2% | 1.00 | 1.00 | 0.92 | 0.89 | 0.95 | 0.96 | 0.94 | 0.96 |
| CD56+ NK | 6.6% | 1.00 | 1.00 | 0.95 | 1.00 | 0.95 | 0.99 | 0.95 | 0.99 |
| DN | 3.5% | 1.00 | 1.00 | 0.81 | 0.69 | 0.90 | 0.85 | 0.83 | 0.76 |
| [not naive CD8+] | 5.4% | 1.00 | 1.00 | 0.90 | 0.85 | 0.94 | 0.92 | 0.93 | 0.91 |
| CD123+ pDC | 0.83% | 1.00 | 1.00 | 0.98 | 1.00 | 0.98 | 0.99 | 0.98 | 0.99 |
| DN DCs | 2.7% | 1.00 | 1.00 | 0.90 | 0.89 | 0.91 | 0.92 | 0.92 | 0.93 |
| DCs- | 2% | 0.99 | 0.99 | 0.37 | 0.23 | 0.20 | 0.10 | | |
| CD1c DCs | 0.37% | 0.99 | 0.99 | 0.56 | 0.42 | 0.43 | 0.30 | 0.57 | 0.44 |
| CD141+DCs | 0.04% | 0.94 | 0.93 | 0.62 | 0.52 | 0.64 | 0.51 | 0.56 | 0.45 |
| **Medians** | | 1.00 | 1.00 | 0.90 | 0.87 | 0.91 | 0.92 | 0.92 | 0.91 |
| **Means** | | 0.99 | 0.99 | **0.83** | **0.79** | 0.81 | 0.78 | **0.72** | **0.69** |

**Table S9. Performance metrics for each population and classifier in the OMIP-044 dataset.**

#### OMIP-047

| Classifier | | MLP | | LDA | | PhenoGraph | | FlowSOM | |
|---|---|---|---|---|---|---|---|---|---|
| Population | Freq-uency | F1-score | CS | F1-score | CS | F1-score | CS | F1-score | CS |
| Naive | 45% | 1.00 | 1.00 | 0.96 | 0.97 | 0.94 | 0.93 | 0.93 | 0.92 |
| MZ B cells | 35% | 1.00 | 1.00 | 0.94 | 0.93 | 0.92 | 0.88 | 0.90 | 0.85 |
| IgA, FSC-A Neg IgA IgG1 IgG3 | 7.3% | 0.99 | 0.99 | 0.85 | 0.81 | 0.84 | 0.73 | 0.61 | 0.50 |
| IgA, FSC-A IgA+ | 5.5% | 0.99 | 0.99 | 0.90 | 0.89 | 0.90 | 0.89 | | |
| IgG1+ | 3.6% | 0.98 | 0.99 | 0.86 | 0.84 | 0.88 | 0.87 | 0.87 | 0.86 |
| Transitional | 2.2% | 0.98 | 0.98 | 0.86 | 0.83 | 0.72 | 0.59 | 0.42 | 0.27 |
| IgG3+ | 0.75% | 0.98 | 0.99 | 0.92 | 0.95 | 0.91 | 0.91 | 0.92 | 0.94 |
| PB | 0.38% | 0.88 | 0.86 | 0.55 | 0.41 | 0.49 | 0.36 | 0.71 | 0.66 |
| Medians | | 0.98 | 0.99 | 0.88 | 0.87 | 0.89 | 0.87 | 0.79 | 0.75 |
| Means | | 0.97 | 0.98 | 0.85 | 0.83 | **0.83** | 0.77 | 0.67 | 0.62 |

**Table S10. Performance metrics for each population and classifier in the OMIP-047 dataset.**

### OMIP-058

| Classifier | | MLP | | LDA | | PhenoGraph | | FlowSOM | |
|---|---|---|---|---|---|---|---|---|---|
| Population | Freq-uency | F1-score | CS | F1-score | CS | F1-score | CS | F1-score | CS |
| CD4+ Naive T cells CD27+ CD28+ | 13% | 1.00 | 1.00 | 0.93 | 0.91 | 0.94 | 0.94 | 0.94 | 0.94 |
| HLA-DR+ | 14% | 1.00 | 1.00 | 0.97 | 0.98 | 0.97 | 0.98 | 0.97 | 0.98 |
| CD4+ TCM T cells CD27+ CD28+ | 34% | 1.00 | 0.99 | 0.93 | 0.91 | 0.87 | 0.81 | 0.87 | 0.82 |
| CD8+ TEMRA T cells CD27- CD28- | 9% | 0.99 | 0.99 | 0.76 | 0.64 | 0.93 | 0.95 | 0.94 | 0.96 |
| CD4+ TCM T cells CD27- CD28+ | 3.5% | 0.99 | 0.99 | 0.81 | 0.73 | 0.73 | 0.63 | 0.70 | 0.57 |
| CD8+ TEM T cells CD27+ CD28+ | 5.4% | 0.99 | 0.99 | 0.84 | 0.79 | 0.72 | 0.62 | 0.76 | 0.69 |
| CD4+ TEM T cells CD27+ CD28+ | 3.3% | 0.99 | 0.98 | 0.64 | 0.49 | | | | |
| CD8+ TCM T cells CD27+ CD28+ | 1.1% | 0.99 | 0.98 | 0.66 | 0.52 | | | 0.06 | 0.03 |
| CD8+ Naive T cells CD27+ CD28+ | 1.1% | 0.99 | 0.98 | 0.67 | 0.52 | 0.59 | 0.42 | 0.59 | 0.42 |
| NK CD16- CD56- | 2.4% | 0.98 | 0.99 | 0.78 | 0.74 | 0.82 | 0.83 | 0.81 | 0.82 |
| CD8+ TEMRA T cells CD27+ CD28- | 1.2% | 0.98 | 0.98 | 0.52 | 0.40 | 0.69 | 0.59 | 0.61 | 0.52 |
| CD4+ TEM T cells CD27- CD28+ | 1.1% | 0.98 | 0.97 | 0.55 | 0.39 | 0.03 | < 0.01 | | |
| CD4+ TEM T cells CD27- CD28- | 1% | 0.98 | 0.98 | 0.43 | 0.28 | 0.48 | 0.32 | 0.48 | 0.32 |
| Vd1+ Vg9- T cells CD56- | 0.88% | 0.98 | 0.98 | 0.80 | 0.73 | 0.85 | 0.80 | 0.84 | 0.82 |
| CD8+ TEM T cells CD27- CD28- | 0.57% | 0.98 | 0.97 | 0.62 | 0.47 | | | 0.03 | 0.02 |
| CD8+ TEMRA T cells CD27+ CD28+ | 0.55% | 0.97 | 0.96 | 0.62 | 0.49 | 0.08 | 0.04 | 0.55 | 0.42 |
| CD4+ TSCM T cells CD27+ CD28+ | 1% | 0.97 | 0.96 | 0.25 | 0.13 | 0.02 | 0.01 | 0.02 | 0.01 |
| CD8+ TEM T cells CD27- CD28+ | 0.31% | 0.97 | 0.95 | 0.68 | 0.55 | | | 0.69 | 0.59 |
| CD4+ TEMRA T cells CD27- CD28- | 0.18% | 0.96 | 0.96 | 0.28 | 0.20 | 0.42 | 0.27 | 0.45 | 0.30 |
| Vd2 high Vg9+ T cells | 0.68% | 0.96 | 0.97 | 0.95 | 0.95 | 0.95 | 0.94 | 0.94 | 0.93 |
| Mature NK | 2.5% | 0.95 | 0.94 | 0.90 | 0.87 | 0.76 | 0.64 | 0.88 | 0.83 |
| CD8+ TSCM T cells CD27+ CD28+ | 0.14% | 0.95 | 0.92 | 0.33 | 0.21 | | | 0.02 | 0.01 |
| CD4+ TEMRA T cells CD27- CD28+ | 0.03% | 0.95 | 0.93 | 0.28 | 0.17 | | | | |
| CD4+ TEMRA T cells CD27+ CD28+ | 0.12% | 0.94 | 0.92 | 0.36 | 0.21 | | | | |
| CD4+ TEM T cells CD27+ CD28- | 0.07% | 0.94 | 0.90 | 0.40 | 0.28 | | | | |
| Vd1+ Vg9- T cells CD56+ | 0.12% | 0.94 | 0.91 | 0.63 | 0.55 | | | | |
| MAIT cells | 0.51% | 0.93 | 0.92 | 0.72 | 0.61 | 0.90 | 0.87 | 0.90 | 0.88 |
| Q2: Vd1+ Vg9+ | 0.08% | 0.93 | 0.95 | 0.75 | 0.72 | 0.81 | 0.81 | 0.78 | 0.76 |
| iNKT cells | 0.07% | 0.93 | 0.92 | 0.93 | 0.90 | 0.92 | 0.92 | 0.96 | 0.97 |
| CD8+ TCM T cells CD27- CD28+ | 0.06% | 0.92 | 0.89 | 0.48 | 0.34 | | | | |
| Vd2 low Vg9+ T cells CD56- | 0.55% | 0.90 | 0.87 | 0.80 | 0.75 | 0.89 | 0.86 | 0.87 | 0.85 |
| CD4+ TSCM T cells CD27- CD28+ | 0.10% | 0.89 | 0.85 | 0.28 | 0.16 | | | | |
| Vd2 low Vg9+ T cells CD56+ | 0.20% | 0.82 | 0.77 | 0.68 | 0.57 | | | | |
| Terminal NK | 0.21% | 0.77 | 0.62 | 0.56 | 0.43 | | | 0.07 | 0.03 |
| Early NK | 0.92% | 0.73 | 0.64 | 0.82 | 0.78 | 0.85 | 0.80 | 0.84 | 0.80 |
| Medians | | 0.97 | 0.96 | 0.67 | 0.55 | 0.48 | 0.32 | 0.59 | 0.42 |
| Means | | 0.95 | 0.93 | 0.65 | 0.55 | 0.43 | 0.40 | 0.47 | 0.44 |

**Table S11. Performance metrics for each population and classifier in the OMIP-058 dataset.**

### OMIP-069

**Scores in OMIP-069 dataset for 43 populations**

| Classifier | | MLP | | LDA | | PhenoGraph | | FlowSOM | |
|---|---|---|---|---|---|---|---|---|---|
| Population | Freq-uency | F1-score | CS | F1-score | CS | F1-score | CS | F1-score | CS |
| Naive CD4+ | 29% | 0.98 | 0.98 | 0.95 | 0.95 | 0.93 | 0.93 | 0.90 | 0.90 |
| B-2 CD27- | 6.8% | 0.98 | 0.97 | 0.91 | 0.91 | 0.93 | 0.93 | 0.93 | 0.93 |
| Naive CD8+ | 9.6% | 0.97 | 0.97 | 0.90 | 0.91 | 0.96 | 0.97 | 0.96 | 0.97 |
| Mature NK | 9.5% | 0.96 | 0.94 | 0.92 | 0.90 | 0.93 | 0.92 | 0.92 | 0.92 |
| B-1 IgG | 0.97% | 0.95 | 0.94 | 0.94 | 0.93 | 0.92 | 0.91 | 0.92 | 0.92 |
| Early Effector CD4+ | 4.1% | 0.95 | 0.91 | 0.71 | 0.56 | 0.38 | 0.26 | 0.39 | 0.25 |
| Central Memory CD4+ CXCR5-, CXCR3- | 4.9% | 0.94 | 0.91 | 0.74 | 0.63 | 0.50 | 0.37 | 0.52 | 0.37 |
| Terminal Effector CD45RA+ CD8+ | 0.94% | 0.94 | 0.91 | 0.56 | 0.45 | 0.28 | 0.18 | 0.31 | 0.20 |
| Early like effector CD4+ | 2.2% | 0.94 | 0.89 | 0.61 | 0.46 | | | | |
| Central Memory CD4+ CXCR5-, CXCR3 | 4.6% | 0.93 | 0.90 | 0.67 | 0.53 | 0.48 | 0.36 | 0.51 | 0.37 |
| CD2+ CD8+ NKT | 1.3% | 0.92 | 0.87 | 0.75 | 0.66 | 0.47 | 0.33 | 0.48 | 0.35 |
| Terminal NK | 0.59% | 0.92 | 0.89 | 0.67 | 0.53 | | | | |
| B-2 CD27+ | 0.68% | 0.92 | 0.90 | 0.51 | 0.37 | 0.51 | 0.37 | 0.46 | 0.34 |
| B-1 IgG- IgM- | 0.73% | 0.91 | 0.92 | 0.86 | 0.85 | 0.89 | 0.90 | 0.84 | 0.82 |
| Central Memory CD4+ CXCR5, CXCR3 | 2.2% | 0.91 | 0.86 | 0.66 | 0.53 | 0.51 | 0.38 | | |
| Early Effector CD8+ | 2.6% | 0.91 | 0.84 | 0.67 | 0.54 | 0.51 | 0.38 | 0.53 | 0.40 |
| Intermediate Effector CD8+ | 0.45% | 0.89 | 0.81 | 0.57 | 0.45 | | | | |
| CD4- CD8- | 0.69% | 0.88 | 0.82 | 0.74 | 0.62 | 0.52 | 0.40 | 0.55 | 0.42 |
| CD123+ pDCs | 0.39% | 0.88 | 0.82 | 0.86 | 0.82 | 0.79 | 0.75 | 0.75 | 0.72 |
| B-1 IgM+ | 0.21% | 0.88 | 0.81 | 0.62 | 0.47 | | | | |
| CCR7- CD45RA+ TCRgD+ | 0.57% | 0.87 | 0.81 | 0.68 | 0.55 | 0.75 | 0.66 | 0.57 | 0.45 |
| Central Memory CD8+ | 0.34% | 0.87 | 0.79 | 0.51 | 0.35 | 0.13 | 0.07 | 0.01 | 0.01 |
| Early NK | 0.22% | 0.87 | 0.81 | 0.67 | 0.56 | 0.31 | 0.15 | 0.81 | 0.73 |
| Terminal Effector | 0.20% | 0.86 | 0.79 | 0.49 | 0.36 | | | | |
| Terminal Effector CD4+ | 0.14% | 0.85 | 0.77 | 0.37 | 0.24 | | | | |
| Early like effector CD8+ | 0.38% | 0.84 | 0.75 | 0.37 | 0.24 | | | 0.06 | 0.03 |
| CD2+ CD8- | 0.22% | 0.83 | 0.79 | 0.51 | 0.40 | | | 0.02 | 0.01 |
| CCR7- CD45RA- TCRgD+ | 0.32% | 0.82 | 0.71 | 0.44 | 0.28 | 0.26 | 0.15 | 0.30 | 0.17 |
| CCR7+ TCRgD+ | 0.12% | 0.77 | 0.74 | 0.41 | 0.28 | | | < 0.01 | < 0.01 |
| ILCs CD4- | 0.43% | 0.75 | 0.63 | 0.17 | 0.09 | 0.04 | 0.02 | 0.33 | 0.22 |
| ILCs CD2+ CD4+ | 0.13% | 0.73 | 0.63 | 0.48 | 0.32 | | | | |
| CD4+ CD8+ T | 0.11% | 0.72 | 0.62 | 0.57 | 0.47 | 0.45 | 0.32 | | |
| Classical monocytes | 12% | 0.69 | 0.54 | 0.72 | 0.57 | 0.73 | 0.60 | 0.73 | 0.59 |
| CD11c+ CD16+ DCs | 0.20% | 0.65 | 0.52 | 0.41 | 0.30 | | | 0.12 | 0.07 |
| ILCs CD2- CD4- | 0.09% | 0.62 | 0.52 | 0.22 | 0.13 | 0.42 | 0.33 | 0.39 | 0.30 |
| CD2- NKT cells | 0.06% | 0.62 | 0.50 | 0.47 | 0.35 | | | | |
| Intermediate monocytes | 1.2% | 0.56 | 0.40 | 0.60 | 0.44 | 0.47 | 0.31 | 0.53 | 0.37 |
| CD1c+ DCs | 0.09% | 0.47 | 0.34 | 0.24 | 0.16 | | | 0.21 | 0.14 |
| Plasmablasts | 0.03% | 0.43 | 0.30 | 0.28 | 0.17 | 0.22 | 0.12 | 0.33 | 0.20 |
| CD1c- CD141- DCs | 0.07% | 0.34 | 0.22 | 0.28 | 0.18 | 0.05 | 0.03 | 0.10 | 0.07 |
| Non classical monocytes | 0.15% | 0.02 | 0.01 | 0.16 | 0.09 | 0.04 | 0.02 | 0.08 | 0.03 |
| CD16- CD14- monocytes | 0.30% | < 0.01 | < 0.01 | 0.04 | 0.02 | 0.16 | 0.10 | 0.05 | 0.03 |
| CD1c- CD141+ DCs | 0.01% | | | 0.10 | 0.06 | | | 0.02 | 0.01 |
| **Medians** | | 0.87 | 0.81 | 0.57 | 0.45 | 0.28 | 0.15 | 0.31 | 0.20 |
| **Means** | | 0.76 | 0.70 | 0.56 | 0.46 | 0.34 | 0.28 | 0.34 | 0.28 |

**Table S12. Performance metrics for each population and classifier in the OMIP-069 dataset.**

### OMIP-077

| Classifier | | MLP | | LDA | | PhenoGraph | | FlowSOM | |
|---|---|---|---|---|---|---|---|---|---|
| Population | Freq-uency | F1-score | CS | F1-score | CS | F1-score | CS | F1-score | CS |
| Neutrophils | 90% | 1.00 | 1.00 | 1.00 | 1.00 | 0.99 | 1.00 | 1.00 | 1.00 |
| T cells | 4.8% | 1.00 | 1.00 | 0.99 | 1.00 | 0.99 | 0.98 | 0.99 | 0.98 |
| Monocytes | 3.6% | 0.98 | 0.98 | 0.87 | 0.82 | 0.97 | 0.97 | 0.97 | 0.97 |
| Basophils | 0.28% | 0.98 | 0.98 | 0.94 | 0.97 | 0.97 | 0.96 | 0.97 | 0.96 |
| Non MZB cells | 0.30% | 0.97 | 0.97 | 0.76 | 0.68 | 0.81 | 0.75 | 0.82 | 0.75 |
| MZB cells | 0.34% | 0.96 | 0.96 | 0.76 | 0.67 | | | | |
| Eosinophils | 0.36% | 0.94 | 0.92 | 0.65 | 0.56 | 0.99 | 1.00 | 0.99 | 1.00 |
| NK cells | 0.50% | 0.92 | 0.93 | 0.78 | 0.74 | 0.97 | 0.98 | 0.96 | 0.98 |
| Plasma cells | 0.01% | 0.76 | 0.68 | 0.46 | 0.33 | 0.91 | 0.88 | 0.48 | 0.35 |
| CD1c+ mDCs | 0.02% | 0.72 | 0.63 | 0.22 | 0.14 | | | | |
| CD141- mDCs | 0.05% | 0.72 | 0.62 | 0.13 | 0.08 | 0.01 | < 0.01 | | |
| CD141+ mDCs | 0.04% | 0.71 | 0.61 | 0.02 | 0.01 | | | | |
| Progenitors | 0.03% | 0.68 | 0.50 | 0.25 | 0.15 | 0.91 | 0.98 | 0.06 | 0.03 |
| pDCs | 0.01% | 0.44 | 0.31 | 0.30 | 0.23 | 0.65 | 0.55 | | |
| Medians | | 0.93 | 0.93 | 0.70 | 0.62 | 0.91 | 0.92 | 0.65 | 0.55 |
| Means | | **0.84** | 0.79 | 0.57 | **0.53** | **0.65** | **0.65** | 0.52 | **0.50** |

**Table S13. Performance metrics for each population and classifier in the OMIP-077 dataset.**

### ESHGHI

| Classifier | | MLP | | LDA | | PhenoGraph | | FlowSOM | |
|---|---|---|---|---|---|---|---|---|---|
| Population | Frequency | F1-score | CS | F1-score | CS | F1-score | CS | F1-score | CS |
| CD4 N | 16% | 1.00 | 1.00 | 0.92 | 0.92 | 0.91 | 0.89 | 0.91 | 0.89 |
| Cd14hi Mono | 20% | 0.99 | 1.00 | 0.94 | 0.95 | 0.91 | 0.87 | 0.91 | 0.86 |
| CD16hi NK | 12% | 0.99 | 1.00 | 0.96 | 0.98 | 0.93 | 0.93 | 0.93 | 0.93 |
| CD8 N | 4.7% | 0.99 | 0.99 | 0.90 | 0.88 | 0.93 | 0.92 | 0.93 | 0.92 |
| CD4 CM | 7.3% | 0.99 | 0.99 | 0.85 | 0.77 | 0.09 | 0.04 | 0.15 | 0.07 |
| gd T | 1.9% | 0.99 | 0.99 | 0.94 | 0.94 | 0.89 | 0.85 | 0.89 | 0.85 |
| CD8 EM | 3% | 0.99 | 0.99 | 0.78 | 0.67 | 0.10 | 0.04 | 0.17 | 0.08 |
| CD8 Temra | 5.1% | 0.99 | 0.99 | 0.84 | 0.78 | 0.80 | 0.71 | 0.78 | 0.69 |
| CD4 EM | 10% | 0.99 | 0.99 | 0.88 | 0.81 | 0.80 | 0.68 | 0.78 | 0.66 |
| Treg | 2.3% | 0.98 | 0.98 | 0.74 | 0.63 | 0.73 | 0.60 | 0.18 | 0.08 |
| Neutrophils | 1.6% | 0.98 | 0.98 | 0.94 | 0.93 | 0.96 | 0.95 | 0.96 | 0.95 |
| CD16hi Mono | 2.8% | 0.98 | 0.97 | 0.89 | 0.86 | 0.73 | 0.62 | 0.90 | 0.86 |
| Naive B | 2.1% | 0.98 | 0.98 | 0.87 | 0.84 | | | 0.87 | 0.83 |
| CD8 MAIT | 1.4% | 0.97 | 0.97 | 0.82 | 0.77 | 0.82 | 0.76 | 0.59 | 0.46 |
| CD8 CM | 0.73% | 0.97 | 0.96 | 0.65 | 0.52 | | | | |
| Basophils | 1.7% | 0.97 | 0.96 | 0.86 | 0.80 | 0.96 | 0.97 | 0.97 | 0.98 |
| Int Mono | 1.6% | 0.97 | 0.95 | 0.67 | 0.56 | 0.04 | 0.01 | 0.72 | 0.61 |
| HLADRio Mono | 0.71% | 0.96 | 0.95 | 0.69 | 0.55 | | | 0.03 | 0.01 |
| CD4 Temra | 1.5% | 0.96 | 0.95 | 0.47 | 0.32 | 0.02 | 0.01 | 0.02 | < 0.01 |
| CD56hi NK | 0.76% | 0.95 | 0.93 | 0.77 | 0.68 | 0.79 | 0.71 | 0.78 | 0.70 |
| Trans B | 0.68% | 0.95 | 0.94 | 0.78 | 0.72 | 0.67 | 0.54 | 0.87 | 0.85 |
| pDCs | 0.39% | 0.94 | 0.91 | 0.79 | 0.72 | 0.93 | 0.92 | 0.93 | 0.92 |
| Eosinophils | 0.21% | 0.94 | 0.94 | 0.90 | 0.87 | 0.74 | 0.69 | 0.75 | 0.69 |
| Mem B | 0.38% | 0.94 | 0.91 | 0.74 | 0.65 | 0.84 | 0.80 | 0.86 | 0.83 |
| mDCs | 0.35% | 0.93 | 0.89 | 0.49 | 0.36 | 0.55 | 0.42 | 0.56 | 0.44 |
| Plasmablasts | 0.19% | 0.91 | 0.86 | 0.56 | 0.47 | 0.69 | 0.61 | 0.68 | 0.59 |
| Lineage Negative | 0.35% | 0.90 | 0.85 | 0.54 | 0.40 | 0.27 | 0.18 | 0.33 | 0.22 |
| IgD- CD27- B | 0.13% | 0.87 | 0.82 | 0.51 | 0.41 | | | 0.40 | 0.27 |
| Medians | | 0.97 | 0.96 | 0.80 | 0.74 | 0.74 | 0.65 | 0.78 | 0.69 |
| Means | | 0.96 | 0.95 | 0.78 | 0.71 | **0.57** | 0.53 | 0.64 | 0.57 |

**Table S14. Performance metrics for each population and classifier in the ESHGHI dataset.**

GHOSN

| Classifier | | MLP | | LDA | | PhenoGraph | | FlowSOM | |
|---|---|---|---|---|---|---|---|---|---|
| Population | Freq-uency | F1-score | CS | F1-score | CS | F1-score | CS | F1-score | CS |
| B-1 | 22% | 0.99 | 1.00 | 0.97 | 0.99 | 0.95 | 0.94 | 0.90 | 0.89 |
| Large macrophages | 32% | 0.99 | 1.00 | 0.98 | 1.00 | 0.98 | 1.00 | 0.99 | 1.00 |
| B-2 IgM medium | 7.2% | 0.99 | 0.99 | 0.89 | 0.84 | 0.84 | 0.83 | 0.79 | 0.74 |
| T cells | 6.9% | 0.99 | 1.00 | 0.97 | 0.99 | 0.96 | 0.98 | 0.97 | 0.99 |
| Eosinophils | 8.6% | 0.99 | 1.00 | 0.98 | 1.00 | 0.98 | 1.00 | 0.94 | 0.99 |
| B-2 IgM high | 5.3% | 0.98 | 0.98 | 0.82 | 0.76 | 0.75 | 0.69 | 0.67 | 0.56 |
| NK, NKT | 8.3% | 0.97 | 0.98 | 0.84 | 0.84 | 0.89 | 0.93 | 0.89 | 0.92 |
| CD11b+ dendritic | 1.3% | 0.95 | 0.96 | 0.83 | 0.75 | | | 0.86 | 0.81 |
| Developing B | 0.55% | 0.95 | 0.93 | 0.77 | 0.66 | | | 0.83 | 0.75 |
| Small macrophages | 4.6% | 0.94 | 0.91 | 0.94 | 0.95 | 0.81 | 0.74 | 0.92 | 0.91 |
| CD11b- dendritic | 0.23% | 0.93 | 0.93 | 0.63 | 0.53 | 0.77 | 0.69 | 0.83 | 0.80 |
| ckit+ mast cells | 1.8% | 0.90 | 0.87 | 0.68 | 0.56 | 0.69 | 0.56 | 0.66 | 0.54 |
| Neutrophils | 0.36% | 0.84 | 0.80 | 0.39 | 0.27 | 0.43 | 0.28 | 0.01 | 0.01 |
| **Medians** | | 0.97 | 0.98 | 0.84 | 0.84 | 0.81 | 0.74 | 0.86 | 0.81 |
| **Means** | | 0.95 | 0.95 | 0.82 | **0.78** | 0.70 | **0.66** | **0.79** | **0.76** |

**Table S15. Performance metrics for each population and classifier in the GHOSN dataset.**

### LEIPOLD

| Classifier | | MLP | | LDA | | PhenoGraph | | FlowSOM | |
|---|---|---|---|---|---|---|---|---|---|
| Population | Freq-uency | F1-score | CS | F1-score | CS | F1-score | CS | F1-score | CS |
| CD4 T cells CD45RA- , CD27+ | 12% | 1.00 | 1.00 | 0.92 | 0.90 | 0.94 | 0.93 | 0.79 | 0.72 |
| CD4 T cells CD45RA+ , CD27+ | 16% | 0.99 | 1.00 | 0.91 | 0.92 | 0.92 | 0.95 | 0.89 | 0.90 |
| CD8 T cells CD45RA+ , CD27- | 9.1% | 0.99 | 0.99 | 0.84 | 0.78 | 0.67 | 0.55 | 0.69 | 0.59 |
| CD4 T cells CD45RA- , CD27- | 3.4% | 0.99 | 0.99 | 0.87 | 0.81 | 0.85 | 0.80 | | |
| NK cells | 12% | 0.99 | 0.99 | 0.86 | 0.83 | 0.87 | 0.86 | 0.90 | 0.89 |
| CD4- CD8- | 5% | 0.99 | 0.99 | 0.76 | 0.69 | 0.79 | 0.70 | 0.69 | 0.60 |
| CD8 T cells CD45RA- , CD27+ | 6.3% | 0.99 | 0.98 | 0.86 | 0.81 | 0.80 | 0.73 | 0.71 | 0.60 |
| CD8 T cells CD45RA- , CD27- | 2.8% | 0.98 | 0.97 | 0.68 | 0.56 | | | 0.11 | 0.06 |
| CD8 T cells CD45RA+ , CD27+ | 11% | 0.98 | 0.98 | 0.86 | 0.84 | 0.68 | 0.57 | 0.64 | 0.55 |
| CD38 CD56- B cells | 6.2% | 0.98 | 0.98 | 0.76 | 0.65 | 0.39 | 0.21 | 0.13 | 0.04 |
| CD38+CD56 B cells | 4.7% | 0.98 | 0.97 | 0.73 | 0.67 | 0.66 | 0.55 | 0.64 | 0.52 |
| CD16- monocytes | 7.9% | 0.98 | 0.98 | 0.90 | 0.88 | 0.90 | 0.90 | 0.89 | 0.88 |
| pDCs | 0.37% | 0.97 | 0.98 | 0.90 | 0.90 | 0.92 | 0.95 | 0.93 | 0.97 |
| CD4 T cells CD45RA+ , CD27- | 1.1% | 0.97 | 0.95 | 0.30 | 0.20 | 0.06 | 0.02 | 0.03 | 0.02 |
| Basophils | 0.69% | 0.94 | 0.89 | 0.11 | 0.05 | 0.14 | 0.05 | 0.07 | 0.02 |
| mDCs | 1.4% | 0.88 | 0.79 | 0.47 | 0.31 | 0.19 | 0.09 | 0.40 | 0.28 |
| CD56 bright NK cells | 0.44% | 0.70 | 0.56 | 0.70 | 0.62 | 0.73 | 0.64 | 0.72 | 0.63 |
| CD16+ monocytes | 0.04% | 0.70 | 0.55 | 0.05 | 0.03 | 0.22 | 0.14 | 0.06 | 0.03 |
| Medians | | 0.98 | 0.98 | 0.80 | 0.74 | 0.70 | 0.61 | 0.66 | 0.56 |
| Means | | 0.94 | **0.92** | 0.69 | 0.64 | 0.60 | **0.54** | 0.52 | 0.46 |

**Table S16. Performance metrics for each population and classifier in the LEIPOLD dataset.**

PANORAMA

| Population | Frequency | MLP F1-score | MLP CS | LDA F1-score | LDA CS | PhenoGraph F1-score | PhenoGraph CS | FlowSOM F1-score | FlowSOM CS |
|---|---|---|---|---|---|---|---|---|---|
| Eosinophils | 8.8% | 0.99 | 0.99 | 0.82 | 0.73 | 0.73 | 0.61 | 0.66 | 0.53 |
| pDCs | 5.2% | 0.99 | 0.99 | 0.93 | 0.88 | 0.85 | 0.77 | 0.91 | 0.86 |
| Classical Monocytes | 21% | 0.99 | 0.99 | 0.88 | 0.83 | 0.85 | 0.79 | 0.82 | 0.75 |
| CD8 T cells | 3% | 0.98 | 0.98 | 0.90 | 0.88 | 0.50 | 0.37 | 0.77 | 0.68 |
| NKT cells | 1.9% | 0.98 | 0.97 | 0.68 | 0.56 | | | 0.64 | 0.52 |
| Intermediate Monocytes | 7.7% | 0.97 | 0.96 | 0.81 | 0.73 | 0.72 | 0.60 | 0.77 | 0.67 |
| Non classical Monocytes | 1.4% | 0.96 | 0.96 | 0.86 | 0.84 | 0.90 | 0.88 | 0.91 | 0.89 |
| CD4 T cells | 1.6% | 0.96 | 0.97 | 0.81 | 0.77 | 0.80 | 0.74 | 0.79 | 0.72 |
| B cells Frac. D | 3.7% | 0.95 | 0.93 | 0.71 | 0.57 | | | 0.41 | 0.28 |
| NK cells | 1.2% | 0.94 | 0.93 | 0.59 | 0.45 | 0.28 | 0.17 | 0.34 | 0.22 |
| B cells Frac A-C | 1.9% | 0.93 | 0.91 | 0.41 | 0.28 | | | 0.05 | 0.02 |
| Plasma Cells | 0.24% | 0.93 | 0.90 | 0.90 | 0.85 | 0.59 | 0.36 | 0.71 | 0.54 |
| B cells Frac. F | 25% | 0.92 | 0.87 | 0.91 | 0.85 | 0.89 | 0.82 | 0.87 | 0.78 |
| Basophils | 0.57% | 0.92 | 0.88 | 0.55 | 0.39 | | | 0.46 | 0.31 |
| Macrophages | 1% | 0.91 | 0.88 | 0.64 | 0.50 | 0.53 | 0.37 | 0.56 | 0.42 |
| mDCs | 0.30% | 0.87 | 0.82 | 0.64 | 0.50 | 0.28 | 0.13 | 0.60 | 0.48 |
| B cells Frac. E | 9.8% | 0.82 | 0.74 | 0.65 | 0.52 | 0.35 | 0.22 | 0.37 | 0.25 |
| gd T cells | 0.21% | 0.80 | 0.74 | 0.41 | 0.30 | | | 0.01 | 0.01 |
| GMP | 2.1% | 0.69 | 0.54 | 0.50 | 0.35 | 0.25 | 0.14 | 0.43 | 0.30 |
| MEP | 1.8% | 0.60 | 0.45 | 0.39 | 0.26 | 0.36 | 0.22 | 0.36 | 0.23 |
| CMP | 1% | 0.49 | 0.35 | 0.39 | 0.27 | 0.24 | 0.15 | 0.25 | 0.17 |
| MPP | 0.35% | 0.47 | 0.32 | 0.26 | 0.16 | | | | |
| CLP | 0.24% | 0.34 | 0.24 | 0.27 | 0.18 | | | 0.11 | 0.07 |
| HSC | 0.02% | | | | | | | 0.01 | < 0.01 |
| **Medians** | | 0.93 | 0.89 | 0.65 | 0.51 | 0.32 | 0.19 | 0.51 | 0.37 |
| **Means** | | 0.81 | 0.76 | 0.62 | *0.53* | ***0.38*** | *0.31* | 0.49 | *0.40* |

**Table S17. Performance metrics for each population and classifier in the PANORAMA dataset.**

### Section S10: Reproducing tables and figures

To view the tables and figures *interactively* for this final round of testing

- Open MATLAB R2021a or later.
- Type the command `fjb` to start FlowJoBridge.
- Type the command `ClassificationTable.See('dataset name')` where 'dataset name' is 'OMIP-044' or whatever name appears in this document above the tables. The command to see all 9 datasets in this paper needs no parameters: `ClassificationTable.See`
- The first time you do this select the first option "Google Cloud (our demos file)". After that select the last option "My *last* choice".
- Click OK on this dialog box.

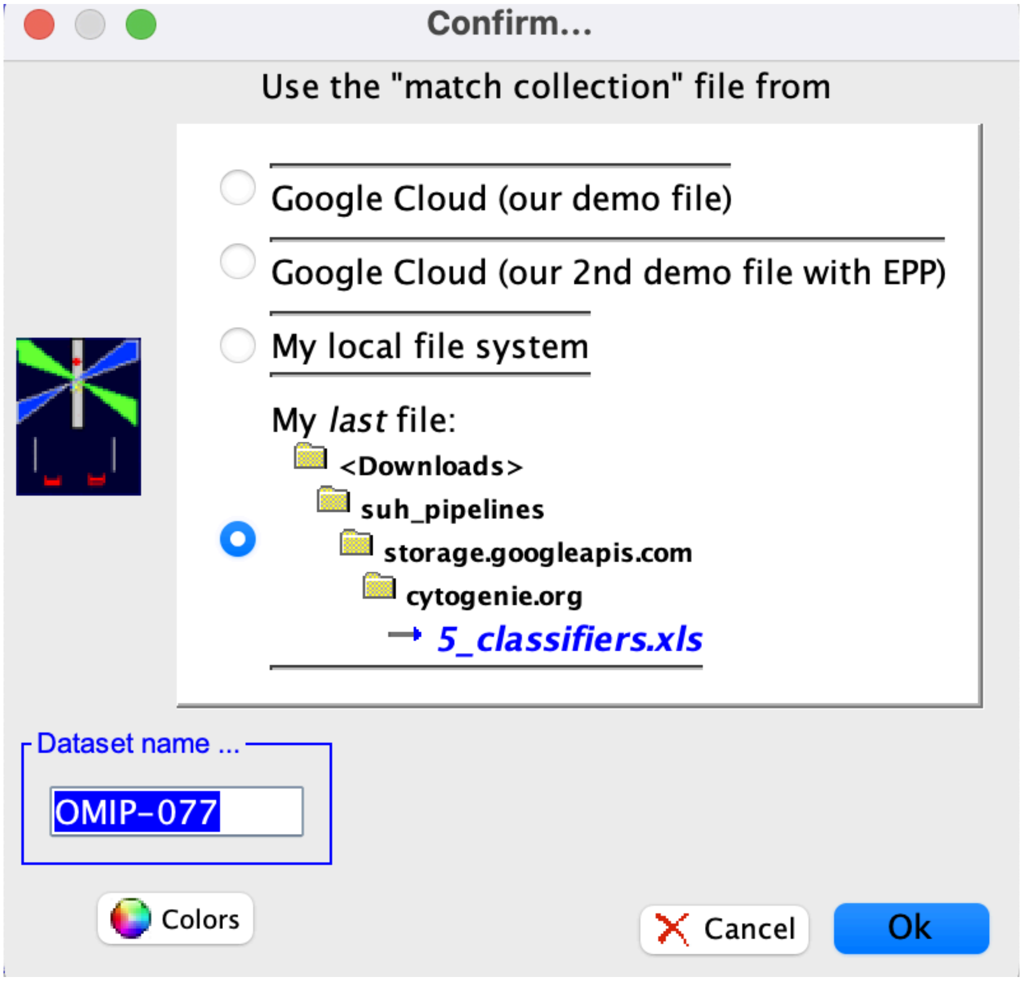

**Figure S5. Dialog box for opening the OMIP-077 tables and figures.**

The first time you run this command download our xls file and colors from the Google cloud.

To get the xls file select the first item in this dialog.

To get the colors click the color button at the bottom left of the dialog and answer yes to the question.

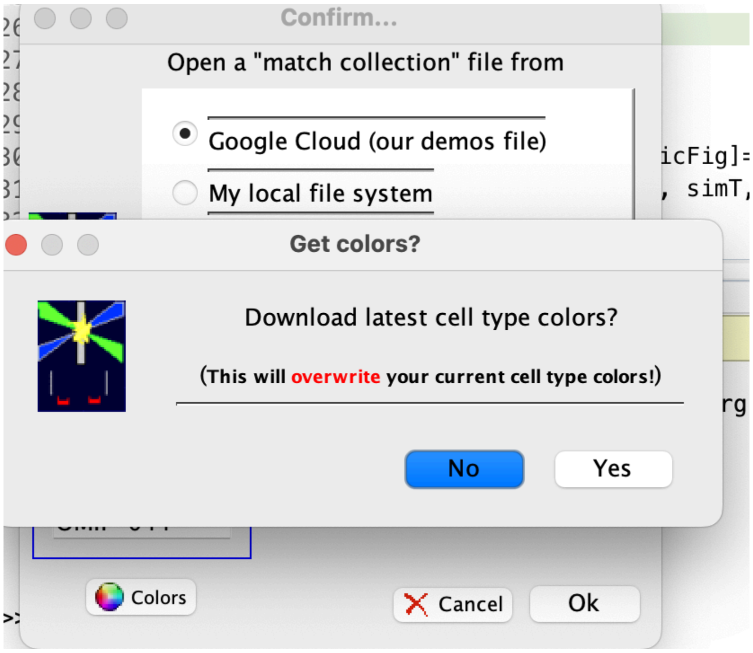

**Figure S6. Dialog box for downloading cell type colors.**

To see table 3 in our paper use the MATLAB command `MlpPaperFunctions.WeakMatches`. To see the box whisker plots and multi-colored tables in section S9 in this document and table 2 in the paper use the MATLAB command `MlpPaperFunctions.Tables`.

For all of this testing and validation we assist our reviewers and readers in reproducing our findings by make everything publicly accessible from our Google Cloud storage:

- All the software we developed.
- All the data sets with which we test it.
- FlowJo workspaces that capture publication recommended manual gating so you don't have to redraw them.
- Documents describing step-by-step how to get the above items and use our FlowJoBridge GUI to get the same results with each gating method and each dataset
